# Ecological opportunity, radiation events and genomic innovations shaped the episodic evolutionary history of papillomaviruses

**DOI:** 10.1101/2020.03.08.982421

**Authors:** Anouk Willemsen, Ignacio G. Bravo

**Affiliations:** Centre National de la Recherche Scientifique (CNRS), Laboratory MIVEGEC (CNRS IRD Univ. Montpellier), Montpellier, France; Centre for Microbiology and Environmental Systems Science, University of Vienna, Vienna, Austria; Centre for Research on the Ecology and Evolution of Diseases (CREES), Montpellier, France

**Keywords:** virus evolution, molecular dating, time-dependent rate phenomenon, virus-host co-evolution and co-phylogeny, infection and cancer

## Abstract

Papillomaviruses (PVs) have a wide host range, infecting mammals, birds, turtles, and snakes. The recent discovery of PVs in different fish species allows for a more complete reconstruction of the evolutionary history of this viral family. In this study we perform phylogenetic dating to analyse evolutionary events that occurred during PV evolution, as well as to estimate speciation and evolutionary rates.

We have used four different data sets to explore and correct for potential biases that particular taxa combinations may introduce during molecular time inference. When considering the evolution of substitution rates we observed that short-term rate estimates are much higher than long-term rate estimates, also known as the time-dependent rate phenomenon. When considering the evolution of viral branching events (as a proxy for speciation rates), we show that these have not been constant through time, suggesting the occurrence of distinct evolutionary events such as adaptive radiations and/or changes in the available host niches. In a joint analysis with host speciation rates, we identified at least four different evolutionary periods, suggesting that the evolution of PVs has been multiphasic, and thus refining the previously suggested biphasic evolutionary scenario. Thanks to the discovery of novel PVs in basal hosts and to the implementation of a time-dependent rate model for molecular dating, our results provide new insights into the evolutionary history of PVs. In this updated evolutionary scenario, ecological opportunity appears as one main driving force for the different radiation and key-innovation events we observe.

## Introduction

Papillomaviruses (PVs) are small non-enveloped circular dsDNA viruses with genome size between 6 and 8 kbp. The minimal PV genome consists of an upstream regulatory region (URR), an early gene region encoding for the E1 and E2 proteins, with in most cases the *E4* gene nested within *E2*, and a late gene region encoding for the L2 and L1 capsid proteins (García-Vallvé et al., 2005). Proteins in the early region are involved in viral replication and cell transformation, while the capsid proteins self assemble to yield virions and encapsidate the genome. During PV evolution, at different time points and in different linages, the viral genomes have acquired, and in cases subsequently lost, the *E5*, *E6* and *E7* oncogenes (Van Doorslaer & McBride, 2016; Willemsen et al., 2019; Willemsen & Bravo, 2019). In chronic infections by certain viruses and in certain hosts, these genes are directly involved in the onset of cancer and behave thus as oncogenes.

PVs have been mainly described to infect mammals, but have also been found in reptiles and birds. More recently, PVs have also been discovered in fish, firstly isolated from a gilt-head sea bream (López-Bueno et al., 2016). Other PV genomes were later isolated from a rainbow trout, two haddocks, and a red snapper (Tisza et al., 2020). This discovery challenged our perspective on the origin of this viral family. Phylogenetic dating studies including this single fish PV genome date back the time to the most recent common ancestor (tmrca) of PVs to 481 (95% HPD: 326─656; Van Doorslaer, Ruoppolo, et al., 2017) and 424 million years ago (95% HPD: 402─446; Willemsen & Bravo, 2019), and thus more ancient than previously thought.

The continuous discovery of novel PVs allows to add pieces to the puzzle of their evolutionary history. Notwithstanding, an enormous bias towards human taxon sampling remains, because certain human PVs (HPVs) are a major public health concern. Indeed, while the majority of PVs cause asymptomatic infections in skin and mucosa, a relatively small number of oncogenic HPVs are associated to malignant lesions that can develop into cancer (Monographs, 2012). For other animals, PVs associated to malignant lesions have been found in a few polyphyletic lineages that mainly infect horses (Scase et al., 2010), cows (Campo, 1997), rabbits (Kreider & Bartlett, 1981) and chamois (Mengual-Chuliá et al., 2014).

Phylogenetic studies have revealed that virus-host codivergence is one of the main driving forces of PV evolution: one third of the viruses’ divergence patterns can be explained by the hosts’ divergence patterns (M. Gottschling et al., 2011). This match allows to identify host divergence times that can be used to date the PV tree (Pimenoff et al., 2017; Rector et al., 2007; Shah et al., 2010). Although virus-host codivergence plays an important evolutionary role in PVs, it is not the only mechanism that has shaped PV diversification. Other evolutionary processes such as recombination (Rector et al., 2008), lineage sorting and host switches have also played a fundamental role in PV evolution, even in recent times (Pimenoff et al., 2017). The best supported scenario proposes a biphasic evolution (Félez-Sánchez et al., 2015; M. Gottschling et al., 2007, 2011), where an early PV radiation would have generated the main extant viral crown groups, followed by independent co-divergence between PVs and their hosts. Consequently, inconsistencies between the virus and host trees are often detected, challenging the inference of ancestral node ages of the PV tree.

Besides node ages, evolutionary rate is one of the key parameters used to characterise the evolutionary history of viruses. Being dsDNA viruses, PVs belong within the group of slow evolving viruses, with rates in the order of 10^−7^ to 10^−9^ nucleotide substitutions per site per year (Pimenoff et al., 2017; Rector et al., 2007; Shah et al., 2010). These values are several orders of magnitudes lower than the rates of their fast evolving counterparts, the RNA viruses (Duffy et al., 2008). However, it is becoming widely accepted that the division between viral types in terms of evolutionary rate is not as strict as historically assumed.

Over the last fifteen years it has become more evident that when performing molecular dating, rate estimates may vary depending on the time frame of measurement, so that rate estimates based on recent calibration nodes are much higher than those based on older calibration nodes (Aiewsakun & Katzourakis, 2015, 2017; Duchene et al., 2014; Ho et al., 2005, 2011). As a consequence, biased divergence times may be inferred, where long-term rates tend to underestimate the divergence time, while short-term rates are prone to overestimation. This time-dependent rate phenomenon has been detected in mitochondria of different organisms (*e.g.* birds, fish, insects, penguins and primates Burridge et al., 2008; García-Moreno, 2004; Ho et al., 2005; Papadopoulou et al., 2010; Subramanian et al., 2009), but has also been communicated for bacteria (Biek et al., 2015; Comas et al., 2013; Feng et al., 2008; Rocha et al., 2006) as well as for multiple levels of viral taxonomy (Aiewsakun & Katzourakis, 2015, 2016; Duchene et al., 2014; Gibbs et al., 2010). Despite the differences in mean substitution rates between large viral groups (*e.g.* between DNA and RNA viruses), the rate decay speed of time-dependent rates is reported to be independent of viral taxonomy (Aiewsakun & Katzourakis, 2016; Duchene et al., 2014; Gibbs et al., 2010). Therefore, corrections proposed to compensate for time-dependent rates have proven to be useful in providing better estimation of the evolutionary time scale of any virus (Aiewsakun & Katzourakis, 2015, 2017; Membrebe et al., 2019).

The time-dependency of molecular rate estimates is a near universal phenomenon and an apparent artefact of the currently available reconstruction approaches. Nonetheless, differences in rate measurements can also reflect biological processes. On the one hand, high values for evolutionary rate, often recovered from short-term rate measurements, are thought to approximate the spontaneous mutation rate (*e.g.* transient deleterious mutations and transient short-sighted adaptations for the current host). On the other hand, low values for evolutionary rate, often recovered from long-term analyses, can better approximate the actual substitution rate (*i.e*. mutations that become fixed) over macroevolutionary timescales (Ho et al., 2011; Simmonds et al., 2019). However, this view does not explain the paradox of why viral genomes are conserved over the long-term while having an apparent unlimited evolutionary potential in terms of population size and mutation rate to evolve and adapt rapidly. Simmonds et al., 2019 proposed an alternative explanatory model where viral genome conservation is best explained by a niche-filling model in which fitness optimization is rapidly achieved in the viral hosts, and therefore viral long-term rates increasingly resemble those of their hosts.

In this study we revisit the evolutionary history of PVs with newly available ancestral fish PV genomes. We selected our calibration points and dated the PV tree using four different data sets based on the consistencies in virus-host codivergence patterns. Two of these data sets contain the virtually complete PV collection hitherto described, while the other two are different versions of reduced data sets to correct for the over-representation of humans and of other economically important hosts. First, we investigate whether a time-dependent rate model better fits PV evolution data than previously used models. Second, we compare evolutionary rates, diversification rates, and node age estimates among the different PV data sets. In addition, we compare the PV evolutionary- and diversification rate estimates with those of the corresponding hosts. Based on our observations we propose an updated evolutionary scenario and point out important events that occurred during the evolution of PVs.

## Materials and Methods

### Data collection and alignments

For this study, 359 full length PV genomes were downloaded from the PaVE (https://pave.niaid.nih.gov/, Van Doorslaer, Li, et al., 2017) and GenBank (https://www.ncbi.nlm.nih.gov/genbank/) databases (Table S1). The *E1*, *E2*, *L2* and *L1* genes were extracted and aligned individually at the amino acid level using MAFFT v.7.271 (Katoh & Standley, 2013), corrected manually, and backtranslated to nucleotides using PAL2NAL v.14 (Suyama et al., 2006). The alignment was filtered using Gblocks v.0.91b (Castresana, 2000), with the following parameters: -t=c, -b1=50% of the number of sequences + 1, -b2=50% of the number of sequences + 1, -b3=8, -b4=3, -b5=a, -b0=3. For tree construction, *E1*, *E2*, *L2* and *L1* were concatenated using a custom perl script.

**Table 1.** Calibration points used for molecular dating of the PV tree. The estimates and the corresponding confidence intervals (CIs) are based on host divergence estimates (TimeTree: http://www.timetree.org/). Numbers in the first column correspond to those used in subsequent graphs and tables in this article to refer to the calibration nodes.

| Number | Calibration node | Divergence estimate<br>(million years ago) | CI<br>(million years) |
| --- | --- | --- | --- |
| 1 | Root | 435 | 425 - 446 |
| 2 | Aves/Testudines | 254 | 240 - 268 |
| 3 | Perissodactyla/Cetartiodactyla | 78 | 74 - 81 |
| 4 | Bovidae1/Cervidae1 | 27.31 | 23.13 - 31.49 |
| 5 | Bovidae2/Cervidae2 | 27.31 | 23.13 - 31.49 |
| 6 | Laurasiatheria/Primates | 96 | 91 - 102 |
| 7 | Platyrrhini/Catarrhini | 43.20 | 40.60 - 45.70 |
| 8 | Homo/Colobus | 29.44 | 27.61 - 31.28 |
| 9 | Homo/Pan | 6.65 | 6.23 - 7.07 |
| 10 | Pan troglodytes/Pan paniscus | 2.82 | 2.40 - 3.24 |
| 11 | Rodentia/Primates | 90 | 85 - 94 |
| 12 | Laurasiatheria/Euarchontoglires | 96 | 91 - 102 |
| 13 | Insectivora/Carnivora | 89 | 83 - 96 |
| 14 | Hyaenidae/Felidae | 40 | 33 - 46 |
| 15 | Primates/Glires | 90 | 85 - 94 |
| 16 | Bovidae3/Cervidae3 | 27.31 | 23.13 - 31.49 |
| 17 | Feliformia/Caniformia | 54 | 52 - 57 |
| 18 | Cricetidae/Muridae | 32.70 | 27.90 - 37.40 |

### Prior phylogenetic analyses

To detect recombinant taxa as well as incongruent taxa between the early gene tree and the late gene tree, we used the filtered concatenated early genes (*E1-E2*) and late genes (*L2-L1*). For these two alignments, the LG+I+Γ protein substitution model was identified as the best suited model using ProtTest v.3.4.2 (Darriba et al., 2011). ML-based phylogenetic analyses were conducted using RAxML v.8.2.9 (Stamatakis, 2014) under the GTR+Γ4 model for the nucleotide alignment using six partitions (three for each gene corresponding to each codon position), under the LG+I+Γ model for the amino acid alignment using two partitions (one for each gene), and using 1000 bootstrap replicates. The trees were rooted using the SaPV1 (*Sparus aurata papillomavirus 1)* sequence (López-Bueno et al., 2016; Fig. S1-S2). Rogue taxa were identified using the algorithm implemented in RAxML (Pattengale et al., 2011). The topologies of the *E1-E2* and *L2-L1* trees were compared using the disagree method in TOPD-fMtS v.3.3 (Puigbo et al., 2007), which allowed us to identify the disagreeing taxa between the two tree topologies.

The previously identified recombinant PVs isolated from Cetaceans (PphPV1-2, TtPV1-7, DdPV1, PsPV1) (Marc Gottschling et al., 2011; Rector et al., 2008; Robles-Sikisaka et al., 2012) also showed to confidently disagree in position between the early and late gene trees in our analyses. Therefore, these recombinant PVs were removed for generating the Full Data set (FD), leaving us with a data set of 343 PV genomes (of which 200 infect humans). Based on the taxa that agree between the *E1-E2* and *L2-L1* trees, we generated two different versions of reduced data sets containing representatives for each PV species and PV type: Representative Data set 1 (RD1) and Representative Data set 2 (RD2). RD1 and RD2 both contain 130 PVs (of which 48 infect humans), and share 85 terminal taxa. RD1 and RD2 were constructed to compensate for certain over-represented host taxa in the FD. For example, the 200 human PVs and the 21 bovine PVs present in the FD were respectively represented by 48 and 6 terminal taxa in the reduced data sets (see Table S1). For the sake of clarity we would like to stress that classification within *Papillomaviridae* is explicitly based on genetic distance, where each PV type is a unique genomic entity. During the establishment of the boundaries for defining the different taxonomic categories within the *Papillomaviridae* (genera – species – type – variant) it was recognized that distribution of pairwise genetic distances is multimodal (De Villiers et al., 2004). The PV working group within the International Committee on the Taxonomy of Viruses decided then to follow these “natural categories” to delineate the boundaries of the taxonomic levels, leading to an official definition of phylogenetic taxa at higher taxonomic levels, as an ensemble of PV types grouped into genera, species and subfamilies (Bernard et al., 2010; Burk et al., 2013; Bzhalava et al., 2015; De Villiers et al., 2004; Van Doorslaer et al., 2018; see also https://talk.ictvonline.org/taxonomy/).

Our FD, RD1 and RD2 contained one single fish PV genome (SaPV1), but during the study four other fish PV genomes were made available in GenBank (accessions: MH510267, MH616908, MH617143, MH617579; Tisza et al., 2020). As the sequences of these new PVs add relevant information to the basal clade of the tree, these genomes were added to the FD, and this new data set containing 347 PV genomes was named FDF (FD + additional fish PVs).

For all four data sets (FDF, FD, RD1, and RD2), the concatenated *E1-E2-L2-L1* alignments were used to construct ML trees with RAxML v.8.2.9 under the GTR+Γ4 model for the nucleotide alignments, using twelve partitions (three for each gene corresponding to each codon position), under the LG+I+Γ model for the amino acid alignments using four partitions (one for each gene), and in both cases using 1000 bootstrap replicates (Fig. S3-S6).

### Phylogenetic time inference

Based on the ML constructed trees, 18 calibration nodes were selected on subtrees where the *E1-E2* and *L2-L1* trees did not show discrepancies (Fig. S1-S2) and where the host tree matched the PV tree. The host time tree (Fig. S7) was recovered from TimeTree (http://www.timetree.org/; Kumar et al., 2017), based on the list of all known PV host species included in this study. Calibration times were based on host divergence molecular clock estimates collected from TimeTree, integrating the corresponding confidence intervals in the prior (Table 1). The effect of the calibration nodes, and therewith forced clades, on the topology of the tree was validated by constructing ML trees constrained to the calibrations used and subsequent comparison to the corresponding unconstrained tree using a Shimodaira-Hasegawa test (Shimodaira & Hasegawa, 1999) as implemented in RAxML (Stamatakis, 2014). At the nucleotide level, the constrained trees did not test significantly worse than the unconstrained trees (Table S2). At the amino acid level, the constrained FDF and RD2 trees tested significantly worse. The nucleotide-based constrained ML trees were used as a starting trees for time inference at the nucleotide level.

**Table 2.**
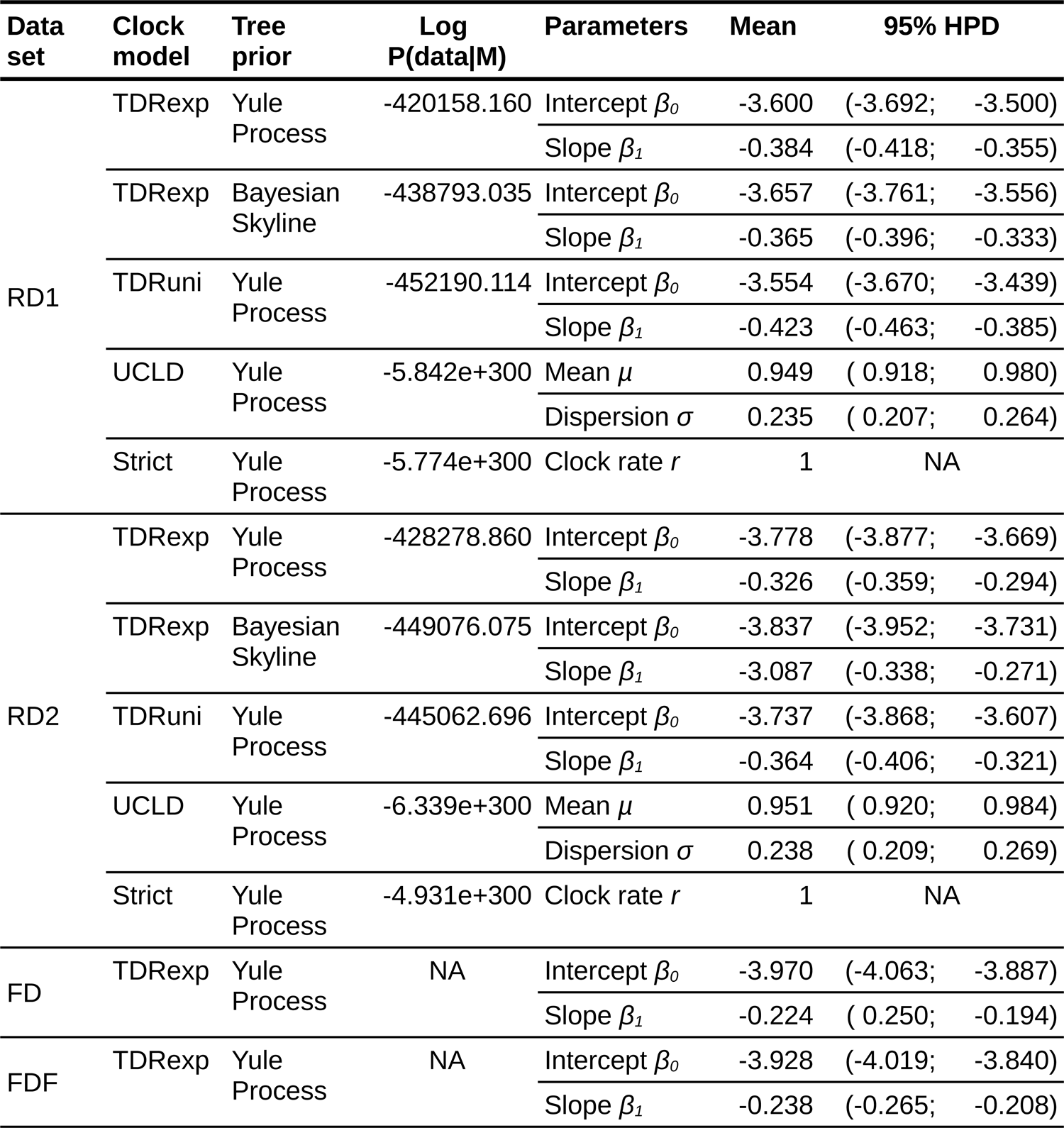
Model fit and regression coefficients for the time-dependent rate clock models. For RD1 and RD2 the model fit of different clock and tree models using (log) marginal likelihood estimates obtained using stepping-stone sampling is given. The exponential time-dependent clock model (TDRexp), the uniform time-dependent clock model (TDRuni), the uncorrelated relaxed clock model (UCLD) and the strict clock mode were compared. The TDRexp model with a Yule tree prior yields the highest log marginal likelihood estimate for both data sets. For RD1, RD2, FD and FDF, the regression coefficients of the TDRexp model were compared. The steeper slopes for RD1 and RD2 indicate a stronger time-dependent rate effect compared to FD and FDF (see also *Fig. 3a*).

Before a time-dependent rate model was implemented in BEAST (Membrebe et al., 2019; Suchard et al., 2018), we manually verified whether a time-dependency of molecular rate estimates exists for PVs. Bayesian time inference was performed at the nucleotide level using BEAST v.1.8.3 (Alexei J Drummond et al., 2012), under the GTR+Γ4 model, using twelve partitions, the uncorrelated relaxed clock model (Alexei J Drummond et al., 2006) with a lognormal distribution and a continuous quantile parametrisation (Li & Drummond, 2012), and the Yule speciation process tree prior (Gernhard, 2008; Yule, 1925). For each calibration node, a prior was defined with a normal distribution around the host divergence estimate and with a standard deviation based on the confidence intervals indicated in Table 1. Time inference was performed separately for each calibration node, on both RD1 and RD2. Thus, per data set, 18 independent time inferences using single calibration nodes were performed. A further description of the manual verification and following correction of time-dependency of molecular rate estimates can be found in Supplementary File 1.

Subsequently, Bayesian time inference was performed at the nucleotide level using BEAST v.1.10.4 (Suchard et al., 2018), under the GTR+Γ4 model, using four partitions (one for each gene). Using data sets RD1 and RD2, four different clock models were tested under the Yule speciation process tree prior (Gernhard, 2008; Yule, 1925): (i) the strict clock model, (ii) the uncorrelated relaxed clock model (Alexei J Drummond et al., 2006) with a lognormal distribution and a continuous quantile parametrisation (UCLD; Li & Drummond, 2012), (iii) a uniform time-dependent rate (TDRuni) clock model, and (iv) an exponential time-dependent rate (TDRexp) clock model (Membrebe et al., 2019). The strict clock model assumes a single substitution rate for the whole tree. The UCLD assumes that that substitution rate along each branch is drawn independently from a lognormal distribution, nonetheless one single mean substitution rate is calculated over the tree. The TDR clock model allows for substitution rate variation over time. The TDRuni and TDRexp epoch models are set up as described in Membrebe et al., 2019, with custom uniform time intervals (with boundaries 0 < 10 < 20 < … < 400 < ∞) and custom exponentially distributed time intervals (with boundaries 0 < 10^-5^ < 10^-4^ < … < 10^2^ < ∞), that are expected to cover the depth of the PV phylogeny. The Yule speciation process branching model assumes a constant speciation rate with no extinction. The clock model with the best model fit was also tested with a coalescent Bayesian Skyline tree prior (A. J. Drummond et al., 2005), that allows the population size to vary stochastically over time.

For each calibration node, a normal distribution was assumed with the host divergence estimate and standard deviation based on the confidence intervals indicated in (Table 1). The standard deviation of the root was relaxed to 50 million years. For each reduced data set, and each model, four independent MCMC chains were run for a maximum of 10^7^ generations, sampling every 10^4^. We compared the model fits of the different clock models with the (log) marginal likelihood estimates obtained using stepping-stone sampling (Baele et al., 2012, 2013), with 100 paths steps, and a chain length of 10^6^. From all models tested with the RD1 and RD2 data sets, the best model was also used for time inference with the FD and FDF data sets. For each of these, seven independent MCMC chains were run for a maximum of 3×10^7^ generations, sampling every 3×10^4^.

### Statistics and graphics

Statistical analyses and graphics were done using R (R Core Team, 2014), with the aid of the packages “ape”, “car”, “dplyr”, “ggfortify”, “ggplot2”, “ggtree”, “lawstat”, “overlapping”, “pgirmess”, “reshape”, “stats”, and “strap”. Computation of optimal breakpoints in the LTT plots was performed using the R package “strucchange”. The optimal number of breakpoints was first calculated using the default ‘strucchange’ parameters. From this analyses the optimal number of breakpoints for the host LTT plot was found to be 3. Since the older nodes contain very few observations (due to limited sampling efforts in basal PVs and thus a limited number of basal branching events), we refined the analyses by specifying a minimal sample size of 3 data points for each segment (parameter h), together with a maximal number of breaks to be calculated set at either 3, 4 or 5. From these analyses, a model with 3 breakpoints (at 148, 64 and 10 Ma) resulted to best fit the phylogenetic reconstruction data of the hosts. The final display of the graphics was designed using Inkscape v.0.92 (https://inkscape.org/en/). The silhouettes in Fig. 1 were obtained from Freepik (https://www.freepik.com/).

**Figure 1.**
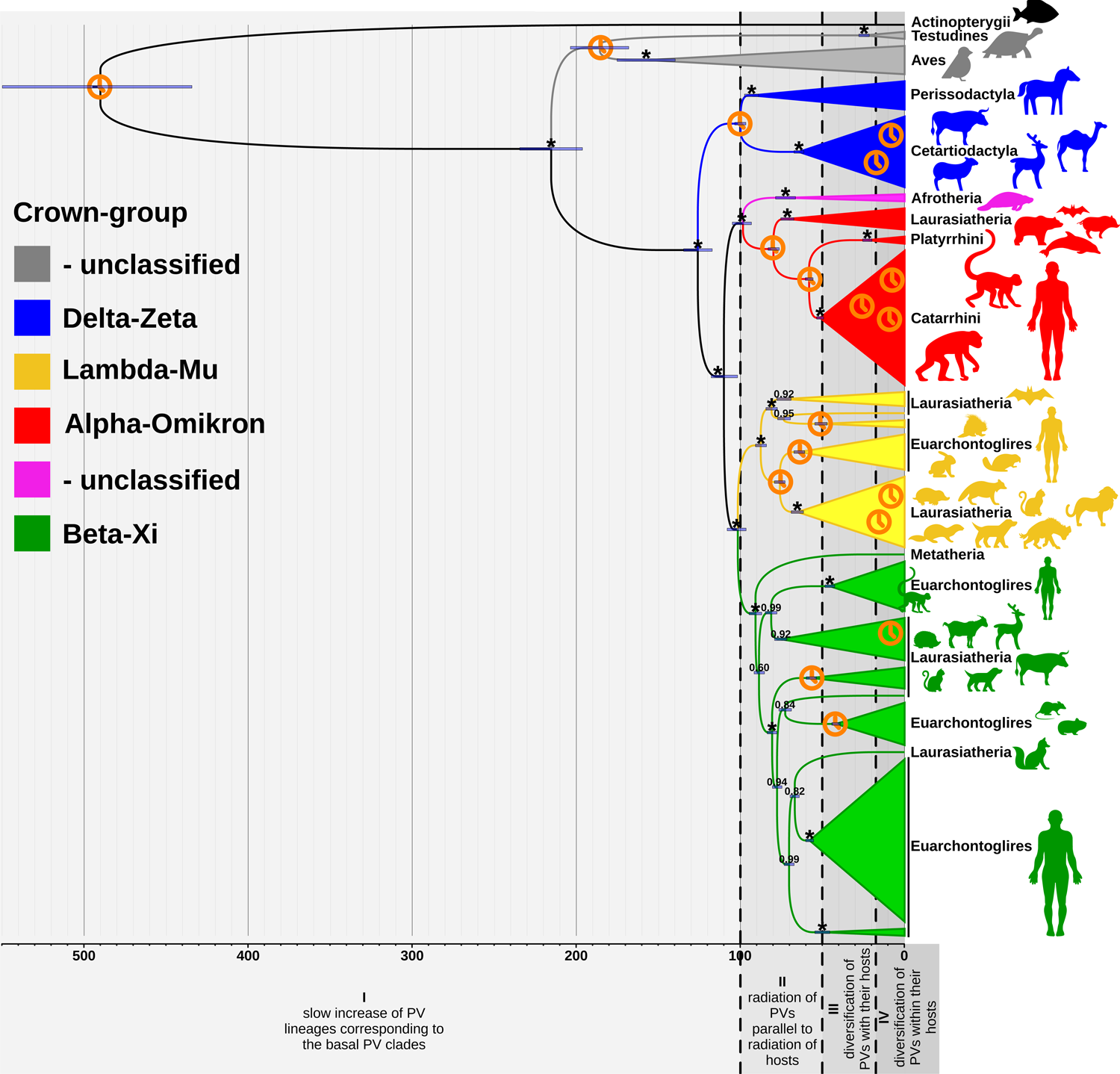
Dated Bayesian phylogenetic tree for reduced data set 2 (RD2) containing 130 PVs. The tree was constructed at the nucleotide level based on the concatenated *E1-E2-L2-L1* genes, using an exponential TDR model and a Yule tree prior. The scale bar is given in million years ago (Ma). Values at the nodes correspond to posterior probabilities, where asterisks indicate full support. Error bars encompass 95% highest posterior density (HPD) intervals for the age of the nodes. Clock symbols indicate the nodes used for calibration. The clades are coloured according to the PV crown group classification, as indicated in the legend on the left. Next to the tree on the right, the taxonomic host group (superorder, superclass, class, order, parvorder, no rank) corresponds to the one in which the corresponding host clades could best be summarized. The cartoon silhouettes illustrate some distinctive members of these clades. The grey shaded areas (delimited by vertical black dashed lines) give an overall overview of four periods that can be distinguished along the evolutionary timeline of PVs, as shown in Fig. 3c.

## Results

### PV crown-groups form well-supported clades, albeit with unclear relative positions among them

In this study we use four different data sets, with a less exhaustive (RD1 and RD2) and a more exhaustive (FD and FDF) representation of all PV genomes in the databases (see Materials and Methods). As some PV lineages ─mainly PVs infecting humans and some economically important species such as cattle, dogs and horses─ are over-represented in the full data sets (FD and FDF), we chose to also work with two different reduced representative data sets (RD1 and RD2). For the construction of RD1 and RD2, the full data sets were used to select one representative for each PV species and type. We would like to stress that for PVs, a type describes a unique genomic entity, genetically different from and within defined boundaries of nucleotide identity to other sister taxa (see also Materials and Methods). RD1 contains representative species with basal and often shorter branches as compared to the derived taxa with often longer branches chosen for RD2. Thus for RD2, more diverse PV genomes are being compared.

For all data sets, the early and the late genes were concatenated (*E1-E2-L2-L1*) and trees were constructed at the nucleotide and amino acid level under a maximum likelihood framework. Overall, we observed well-supported clades for the different PV crown groups (Fig. 1 and Fig. S3-S6): Alpha-OmikronPVs, Beta-XiPVs, Lambda-MuPVs, and Delta-ZetaPVs. Despite the conserved support of these large clades, their relative positions, however, vary for the different trees constructed at the nucleotide and amino acid levels, as well as between the different data sets.

### A time-dependent rate model describes best the evolutionary history of PVs

When performing time inference using one single calibration at a time (Table 1), we observed that a time-dependency of molecular rate estimates exists for PVs, with younger calibrations rendering higher inferred values of the molecular evolutionary rate. As shown in Fig. 2, there is a strong correlation between the substitution rate and the inferred time (Fig. 2a: RD1, *Spearman’s rho* = 0.8225, *S* = 1766, *p* = 2.419e-05; Fig. 2b: RD2, *Spearman’s rho* = 0.5170, *S* = 1470, *p* = 0.0299). For RD1 the substitution rate inferred for the youngest node (2.62 Ma: 1.66×10^-8^ s/s/y) is 3.3 times higher than the substitution rate inferred for the root of the tree (434.11 Ma: 5.03×10^-9^ s/s/y, see also Table S3). For RD2 the substitution rate inferred for the youngest node (3.61 Ma: 0.68×10^-8^ s/s/y) is 1.3 times higher than the substitution rate inferred for the root of the tree (434.19 Ma: 5.22×10^-9^ s/s/y, see also Table S3). The difference in substitution rate for the younger nodes is probably related to the criteria for taxa choice in both reduced data sets, which explore different terminal branch lengths and thus evolutionary distances for recent nodes, with a more limited impact on the values inferred for deeper nodes. When performing time inference using all 18 calibrations nodes together, the variation of evolutionary rates over time is not captured under the uncorrelated relaxed clock model (UCLD). Therefore, we have estimated our node ages by correcting for the time-dependent rate phenomenon by fitting a power-law rate decay model to the single calibration data (as described in Supplementary File 1). Recently, a time-dependent rate (TDR) clock model was implemented in BEAST, that accommodates rate variation through time (Membrebe et al., 2019). When performing time inference using all 18 calibration nodes (Table 1) and the TDR clock model, we observe a strong TDR effect under an epoch structure with both uniformly and exponentially distributed time intervals (Fig. S8). To investigate whether the TDR model indeed better describes the evolutionary history of PVs, we tested the TDR clock model against the UCLD and strict clock model. Both the exponential and uniform TDR clock models yield considerably higher log marginal likelihood estimates compared to the UCLD and strict clock models (Table 2). The exponential TDR model shows the best fit to the data among the clock models being compared. Fig. 3a depicts the molecular rate estimates under the exponential TDR model with a pronounced rate increase towards the present for all four data sets. Comparing all four data sets under the exponential TDR model, we estimate the short-term PV rate to be between 1.29×10^-8^ and 1.42×10^-8^ substitutions/site/year (s/s/y), while the long-term PV rate is estimated to be between 2.42×10^-9^ and 4.58×10^-9^ s/s/y (Table 3). The evolutionary rate estimates for all epochs can be found in Table S4. Under the uniform TDR model and using the reduced data sets, the estimated short-term PV rate (Table S5, between 0-10 Ma: ∼1.39×10^-8^ s/s/y) falls within the range of the rate obtained under the exponential TDR model. The long-term PV rate is estimated to be lower under the uniform TDR model (Table S5, above 400 Ma: ∼1.81×10^-9^ s/s/y), most probably related to the last epoch starting 300 Ma later compared to the last epoch under the exponential TDR model (above 100 Ma). Not unexpected, the estimated short-term PV rates are remarkably higher while the estimated long-term PV rates are lower, than the overall evolutionary rate estimate of ∼8×10^-9^ s/s/y inferred under the UCLD model (Table 3). With the reduced data sets we infer higher short-term and lower long-term rate as compared to the full data sets (Fig. 3a), *i.e.* a stronger time-dependent rate effect, as is evident in the regression coefficients for RD1 and RD2 and the regression coefficients for FD and FDF (Table 2).

**Figure 2.**
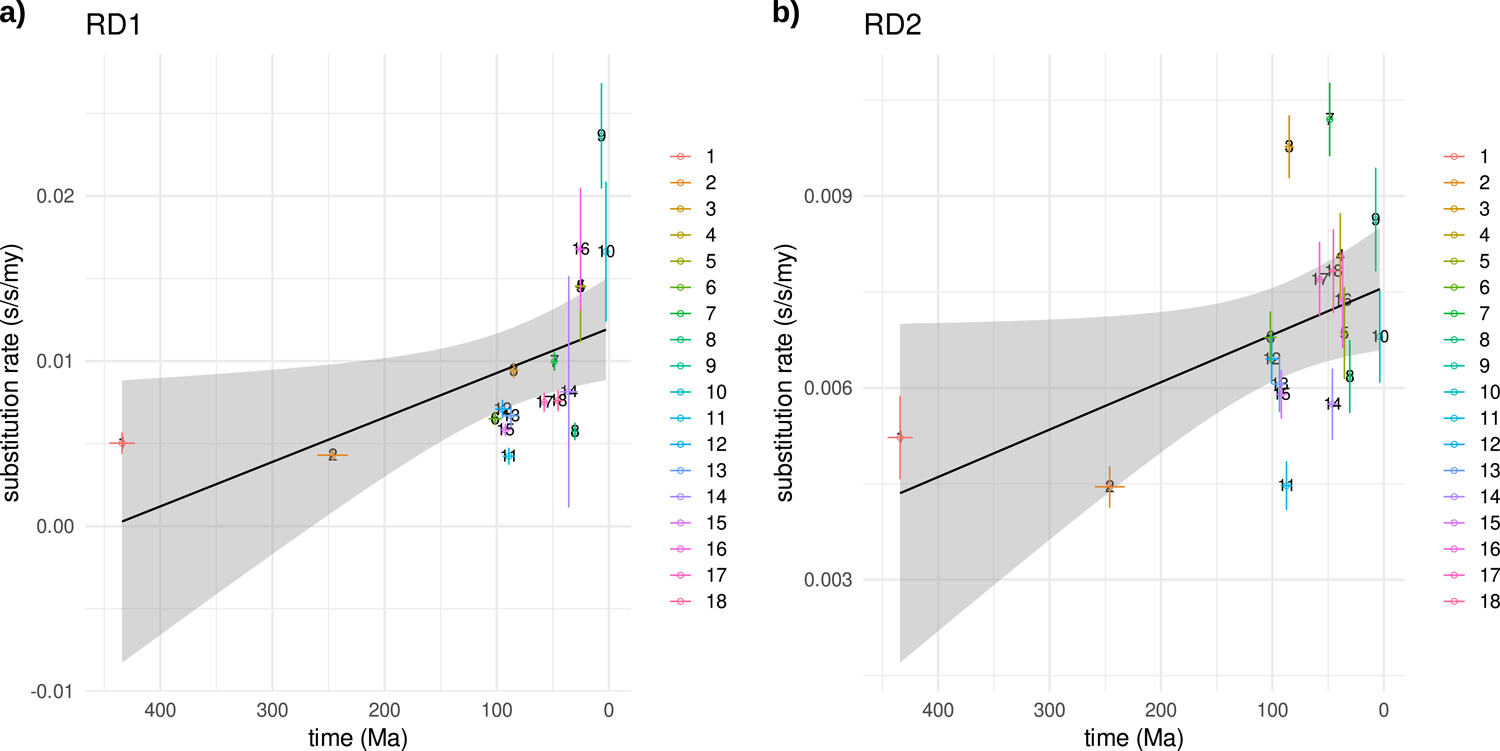
Time-dependent rate phenomenon for PVs. **a)** Results are given for 18 phylogenetic time inferences on RD1. Each inference was performed with a single calibration node, for which the numbers in the legend correspond to those in Table 1. The x-axis represents time from past to present in million years ago (Ma). The y-axis represents the evolutionary rate in substitutions/site/million years (s/s/my). Error bars correspond to the mean and standard deviation of both the inferred node time and the inferred substitution rate. The black line displays the trend of a linear model fit to the data, with the grey shaded area indicating the 95% confidence interval. **b)** The same analysis as in panel a was performed on RD2. Note that the y-axes in panels a and b are not on the same scale.

**Figure 3.**
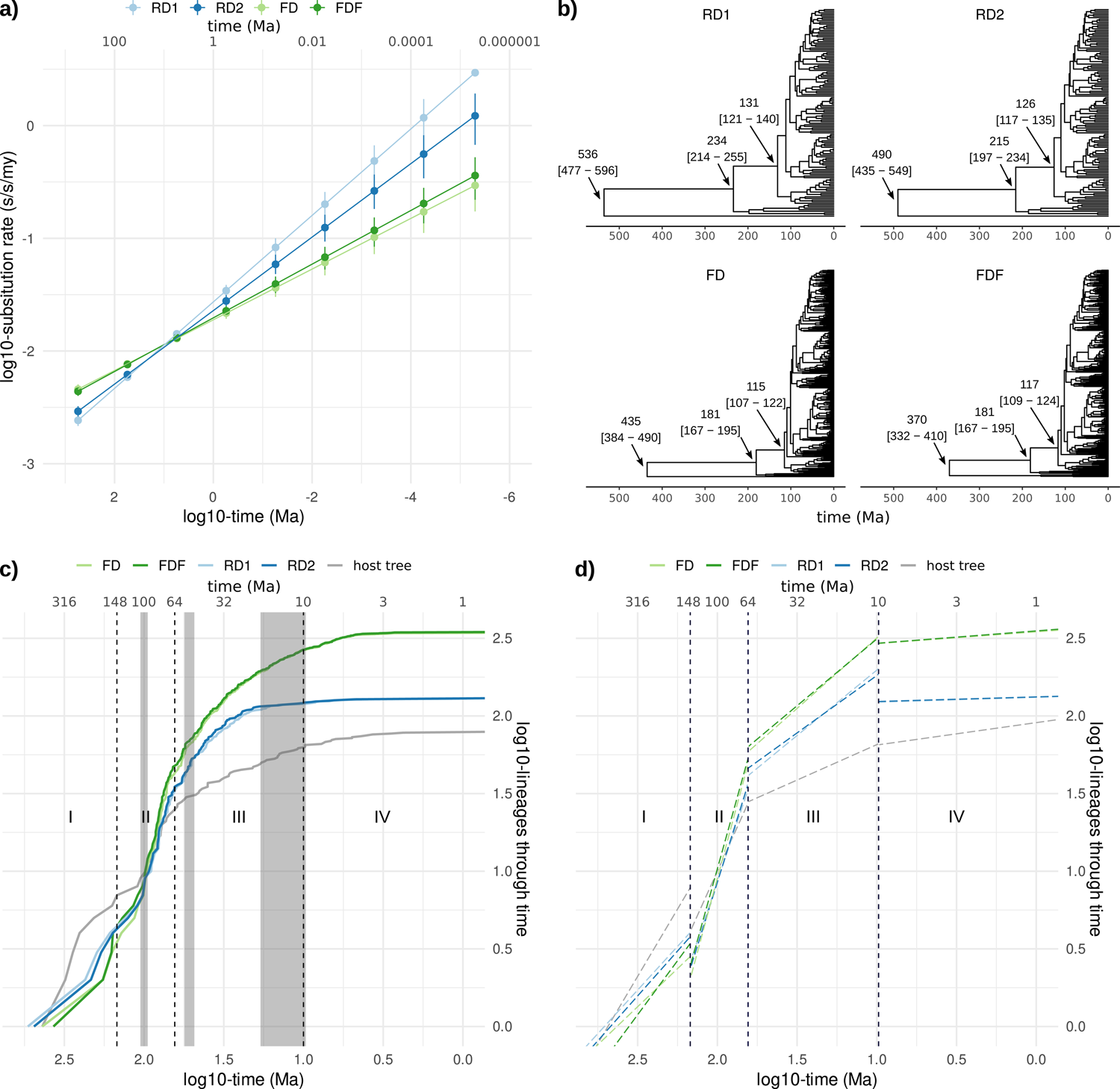
Time-dependent rate epoch modelling and diversification on dated phylogenetic PV trees. Bayesian time inference was performed using two representative reduced data sets (RD1 and RD2) and two full data sets (FD and FDF). **a)** Time-dependent rate effect under an epoch structure with exponentially distributed time intervals. The x-axis represents time from past to present in million years ago (Ma). The y-axis represents the evolutionary rate in substitutions/site/million years (s/s/my). The error bars represent the 95% highest posterior density (HPD) intervals of the rate estimates. Both axes are on a log10 scale. **b)** Dated phylogenetic trees for each of the datasets used, inferred with an exponential TDR model and a Yule tree prior. The x-axis represents time from past to present in Ma. The arrows indicate three basal nodes for which the inferred age is given with the 95% HPD between brackets. These basal nodes are inferred to be much older in RD1 and RD2 when compared to FD and FDF. **c)** Lineages through time (LTT) plot for RD1, RD2, FD, FDF and the host tree. The x-axis represents time from past to present in Ma. The y-axis represents the number of lineages. Both axes are on a log10 scale. The grey vertical bars and black dashed vertical lines separate four evolutionary periods (I-IV) that we observe in the viral and in the host data, respectively. **d)** The slopes of the LTT lines were calculated using the four evolutionary periods of the host data. The slopes are also provided in Table 4. This figure highlights the differences in results obtained for the different data sets, but more importantly, shows that PVs have not diversified at a constant rate.

**Table 3.** The mean, short- and long-term evolutionary rate estimates for PVs. The mean evolutionary rates were inferred by using the uncorrelated relaxed clock model (UCLD) with a Yule tree prior and using all 18 calibration nodes together. The short- and long-term evolutionary rates were inferred by time inference using the exponential TDR model with a Yule tree prior and using all 18 calibration nodes together. Values are in substitutions per site per year (s/s/y).

| Data set | Nr. of taxa | Mean rate (s/s/y) | 95% HPD | Short-term rate (s/s/y) (1 - 10 Ma) | 95% HPD | Long-term rate (s/s/y) (above 100 Ma) | 95% HPD |
| --- | --- | --- | --- | --- | --- | --- | --- |
| RD1 | 130 | $7.90 \times 10^{-9}$ | ($7.45 \times 10^{-9}$; $8.35 \times 10^{-9}$) | $1.42 \times 10^{-8}$ | ($1.27 \times 10^{-8}$; $1.56 \times 10^{-8}$) | $2.42 \times 10^{-9}$ | ($2.16 \times 10^{-9}$; $2.68 \times 10^{-9}$) |
| RD2 | 130 | $8.01 \times 10^{-9}$ | ($7.55 \times 10^{-9}$; $8.47 \times 10^{-9}$) | $1.31 \times 10^{-8}$ | ($1.18 \times 10^{-8}$; $1.45 \times 10^{-8}$) | $2.92 \times 10^{-9}$ | ($2.60 \times 10^{-9}$; $3.24 \times 10^{-9}$) |
| FD | 343 | $7.99 \times 10^{-9}$ | ($7.62 \times 10^{-9}$; $8.36 \times 10^{-9}$) | $1.29 \times 10^{-8}$ | ($1.16 \times 10^{-8}$; $1.41 \times 10^{-8}$) | $4.58 \times 10^{-9}$ | ($4.18 \times 10^{-9}$; $5.06 \times 10^{-9}$) |
| FDF | 347 | $7.68 \times 10^{-9}$ | ($7.31 \times 10^{-9}$; $8.02 \times 10^{-9}$) | $1.31 \times 10^{-8}$ | ($1.19 \times 10^{-8}$; $1.47 \times 10^{-8}$) | $4.39 \times 10^{-9}$ | ($4.00 \times 10^{-9}$; $4.86 \times 10^{-9}$) |

**Table 4.**
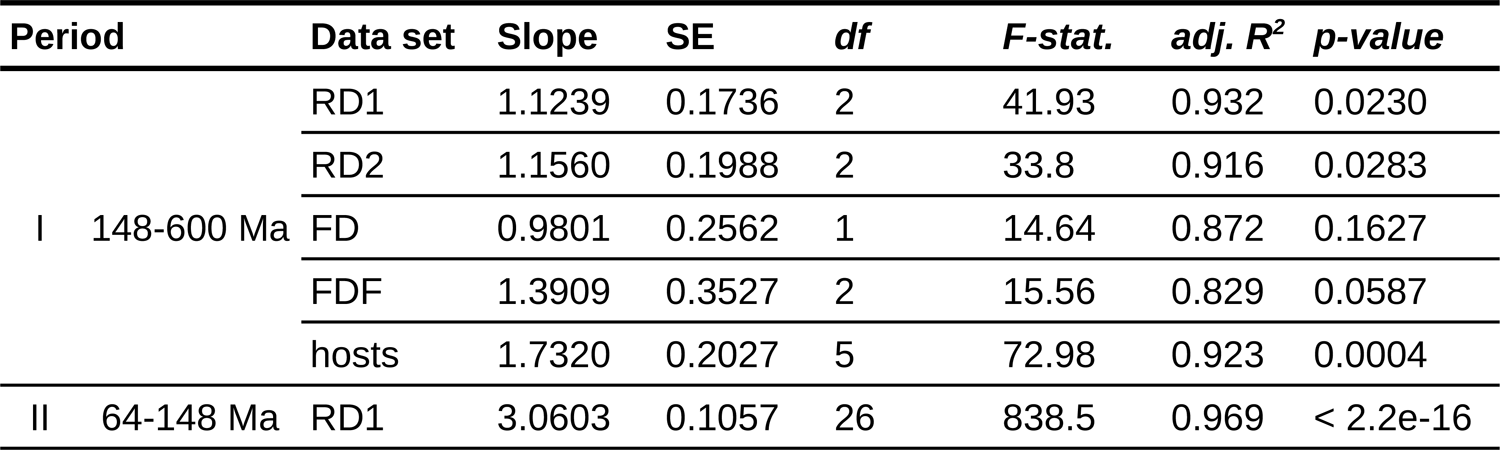
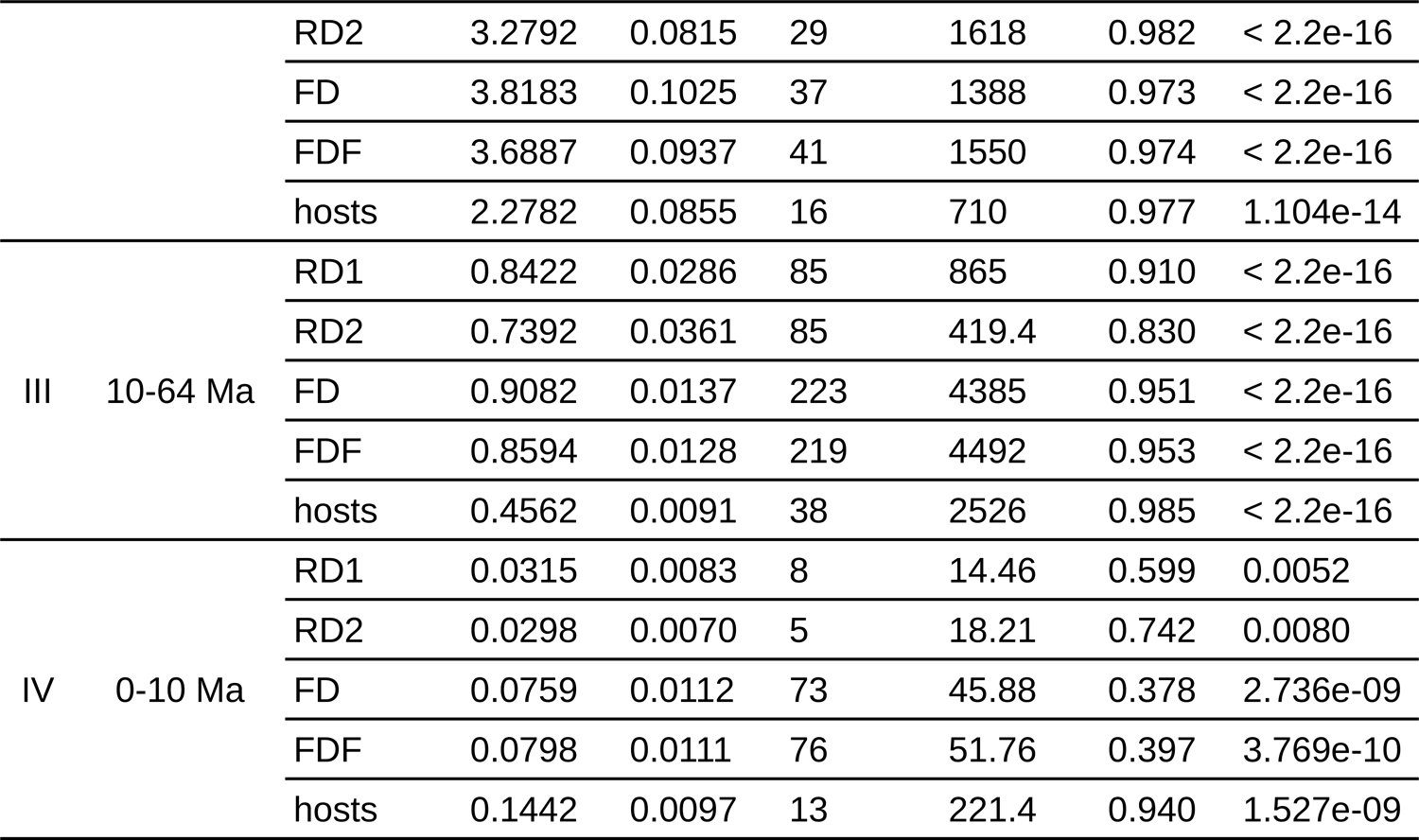
The slopes of the LTT lines in *Fig. 3c*. For each data set (RD1, RD2, FD and FDF) the slope is given for periods I-IV and also shown in *Fig. 3d*. These periods were defined through the computation of breakpoints in regression relationships using the host LTT plot. The slopes are measured in log10 of the number of new lineages per log10 of million years. The standard error (SE), the degrees of freedom (df = N-2), the F-statistic, adjusted R^2^, and the p-value are given to indicate the significance of the calculated slope.

### PVs have not diversified at a constant rate

The dated phylogenetic trees using RD1, RD2, FD, and FDF, with an exponential TDR model and a Yule tree prior are shown in Fig. 3b and Fig. S9-12. A detailed annotated version of the tree obtained using RD2 is depicted in Fig. 1. The main differences between the reduced and full data sets are observed in the ancestral nodes, where those of RD1 and RD2 are inferred to be much older than those of FD and FDF (Fig. 3b). To visualize the timing of PV diversification, we constructed lineages-through-time (LTT) plots based on the dated trees and compared these to the host tree (Fig. 3c). For both viral and host LTT plots we observe different evolutionary periods, where the branching events are not constant through time, suggesting the occurrence of distinct evolutionary events such as adaptive radiations. For viral data we can distinguish four evolutionary periods (divided by grey vertical bars in Fig. 3c), that are based on the observation of breakpoints in the four viral LTT plots (RD1, RD2, FD and FDF). Using the host LTT plot, we computed the optimal breakpoints, allowing us to establish general borders (at 148, 64 and 10 Ma) for four evolutionary periods (divided by black dashed vertical lines in Fig. 3c). Our results are consistent with the main periods proposed for the evolution of the lineage leading to placental mammals: (i) 148 Ma (95% CI: 110.0-158.6) could span the divergence time of Placentals and Marsupials and the initial basal diversification within mammalian orders; (ii) 64 Ma (95% CI: 62.0-66.8) corresponds to intra-ordinal the diversification of placental mammals; and (iii) 10 Ma (95% CI: 7.7-9.7) corresponds to a probable much slower rate of diversification until present. These boundaries inferred based on the host tree roughly match the borders for the observed viral evolutionary periods, and were used to compare the slopes of the viral and host LTT plots (Fig. 3d and Table 4). Our analysis of the LTT slopes indicates that the rate of emergence of novel viral lineages was higher than the rate of novel host lineages in periods II and III (*i.e.* steeper slopes for the emergence of PV lineages). It is important to note that our estimates increase in uncertainty as we approach the root of our tree (period I), essentially due to the limited number of viral taxa retrieved from non-mammalian hosts.

In period I (between ∼600 and ∼148 Ma) we first observe a slow increase of PV lineages that corresponds to the basal PV clades: PVs infecting fish, birds, turtles and ancestral mammals. In period II (between ∼148 and ∼64 Ma), the number of PV lineages surpasses the number of host lineages, growing at a ∼1.5 times faster rate. We interpret that evolutionary period II corresponds to a viral radiation, that goes in parallel with the crown radiation of placental mammals ∼100 Ma, that generated a large number of viral lineages. In period III (between ∼64 and ∼10 Ma) both the PV and host slopes decrease, notwithstanding that the number of PV lineages continues to increase at a rate that is ∼1.8 times faster as compared to the hosts. We interpret that period III corresponds to the diversification of placental mammals within the host crown-groups and the parallel diversification of PVs with their hosts. Lastly, in period IV (between ∼10 Ma and the present) we observe a further decrease in both PV and host slopes. We interpret that in addition to virus host co-evolution, the flattening slopes in periods III and especially IV, indicate a much slower diversification of PVs within their hosts (Fig. 3d and Table 4). The lower slope values for PV diversification in the reduced data sets compared to the full data sets arises most likely from the reduction of the number of terminal taxa associated to individual hosts.

The LTT plots and the LTT slope analysis suggest that the per-lineage speciation rates have not remained constant through time. Thus we can reject the null hypothesis of constant diversification rates on the FDF, FD, RD1, and RD2 data sets, suggesting that the observed trends in PV number of lineages through time is compatible with evolutionary events such as adaptive radiations and/or key adaptations. The Yule tree prior used for time inference assumes a constant speciation rate with no extinction, indicating that this tree prior is an inadequate model. Therefore, we also performed time inference with a Bayesian Skyline tree prior (BS) and the exponential TDR clock model. When comparing the LTT plots of trees inferred with either a Yule or a BS tree prior, we observed that these render highly similar results (Fig. S13). Nevertheless, time inference with a Yule tree prior yields a higher log marginal likelihood estimate than the inference with the BS tree prior (Table 2). This probably indicates that the concatenated PV gene alignments and the calibration nodes are much more informative than the tree prior, which plays thus a minor role on the results.

### Node age estimates on different data sets

For the dated trees constructed on our full and reduced data sets, we obtained the tmrca estimates for different clades along the PV tree (Fig. 4 and Table S6). For those nodes for which the posterior distribution of the age was sampled, we show to what extent the distributions overlap in Fig. S14. Overall, the tmrca estimates obtained by using different data sets are similar. Larger differences are found for the more ancient nodes: the root of the tree, *AvesTestudines/Mammals*, *Aves/Testudines*, *Aves*, and *Mammals* (Fig. 4 and Fig. S14). For these nodes, we obtained older tmrca estimates when using the reduced data sets (RD1 and RD2), as compared to the full data sets (FD and FDF). Due to the scarce number of PVs described for the hosts corresponding to the underlying PV clades, the representative extant taxa are the same in these data sets (see Table S1). Therefore, the ages of these ancient nodes appear to be mostly influenced by the number of mammalian PVs present in the data sets, where removal of overrepresented PVs leads to older time estimates. For the youngest nodes, *Homo/Colobus*, *Homo/Pan* and *Pan troglodytes/Pan paniscus,* inconsistencies in the tmrca estimates were also obtained (Fig. 4 and Fig. S14). For *Homo/Colobus* and *Pan troglodytes/Pan paniscus*, we obtained younger tmrca estimates when using RD1 and RD2, as compared to the FD and FDF, while for *Homo/Pan* this is the inverse.

**Figure 4.**
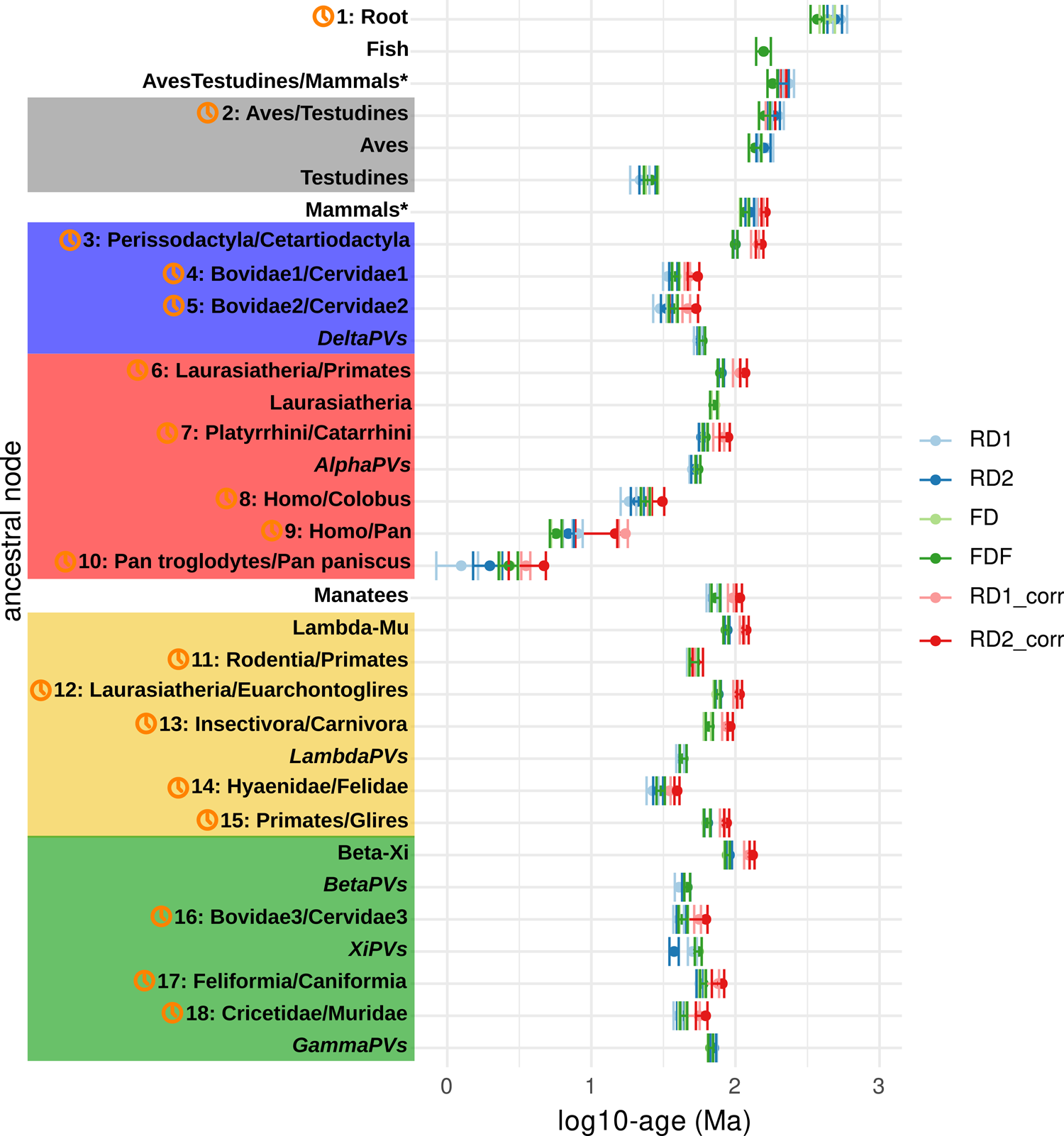
Inferred age for ancestral nodes on dated Bayesian phylogenetic trees. This figure allows to compare for each node the inferred age on the different data sets: RD1, RD2, FD, and FDF. Dated phylogenetic trees were inferred by using the exponential TDR model with a Yule tree prior and 18 calibration nodes. RD1_corr and RD2_corr are corrections for the time-dependent rate phenomenon for trees inferred by time inference using an UCLD model with a Yule tree prior and 18 single calibration nodes (Supplementary File 1). The nodes are ancestors of clades within the PV crowngroups and unclassified clades, matched in colour code with Fig. 1. The nodes used for calibration are indicated with a clock symbol and the corresponding number (see Table 1). The node age is in million years ago (Ma) and on a log10 scale. For RD1, RD2, FD and FDF, error bars encompass 95% highest posterior density (HPD) intervals for the age of the nodes. For RD1_corr and RD2_corr, error bars encompass the lowest and highest inferred 95% confidence intervals (CI) for the 18 different nodes used for error correction. The asterisk indicates that not all the extant PV lineages underlying this node infect mammals as there is one exception in FD and FDF data sets: a PV isolated from a carpet python (MsPV1).

Ultimately, we investigated how the TDRexp clock model performs when compared to corrections for time-dependent rate estimates done by fitting a power-law rate decay model to single calibrations under the UCLD model (Supplementary File 1). The overall comparison for the inferred node ages is shown in Fig. 4. For the *Mammals* node (representing the ancestor of PVs infecting mammals) and for most of the underlying nodes, the corrections estimate the node ages to be older than those inferred by the TDRexp model. This is also the case for the *Homo/Colobus*, *Homo/Pan* and *Pan troglodytes/Pan paniscus* nodes. For the more ancient nodes (*AvesTestudines/Mammals* and *Aves/Testudines*), the corrections estimate the node ages to fall within the range of those inferred by the TDRexp model.

## Discussion

With this study we provide an updated overview on the evolutionary history of PVs, with an emphasis in the expansion and radiation events that have punctuated the timeline of this viral family. Our results show that initial PV evolution seems to be marked by virus-host codivergence, where the number of PV lineages slowly increases during the basal diversification of amniotes. The actual increase is probably much faster than the one we communicate here, because a proper sampling of PVs in basal amniotes and in fishes is still wanting. Subsequently, we observe a viral radiation event that appears to be concomitant with the radiation of the placental mammals. The ecological opportunity for PVs to explore new hosts (vacant niches), can be considered a trigger of the adaptive radiation event we observe. Previous studies have already proposed a link between ecological opportunity and adaptive radiation (Glor, 2010; Yoder et al., 2010). In the case of PVs, adaptive radiation may increase the chances that some of the descendants are better able to exploit the new hosts, thus resulting in species that possess different types of adaptations. When all available niche space becomes filled, we expect rates of lineage accumulation to decrease (Rabosky & Lovette, 2008), and this is exactly what we observe for PVs. After the radiation event, the slope of the LTT plot decreases, and in parallel to virus-host codivergence, independent diversification of PVs occurred. Within the major PV crown groups, the evolution of key innovations allowed PVs to exploit new resources. At the molecular level, the most compelling example is that of *AlphaPVs* infecting humans, where the appearance of the *E5* oncogene (Bravo & Alonso, 2004) triggered an adaptive radiation that generated three viral lineages with different tissue tropisms. Within these three clades, PVs further diversified some evolved the potential to degrade tumor suppressor proteins (Fu et al., 2010; Mesplede et al., 2012; Mirabello et al., 2017; Van Doorslaer et al., 2015; Willemsen et al., 2019).

In this study, we used four different data sets to explore potential biases of the choice of terminal taxa to infer the evolutionary time scale of PVs. The main differences between the full and reduced data sets are apparent in (i) the inferred ages of the basal nodes of the PV tree and (ii) the strength of the time-dependent rate effect. First, the more basal nodes, between roughly 100 and 550 Ma, are generally inferred to be older when using the reduced representative sets of taxa compared to using all taxa (Fig. 3b, Fig. 4 and Fig. S14). Secondly, the time-dependent rate effect is stronger when using the reduced data sets compared to using the full data sets, as evidenced by a steeper slope in Fig. 3a.

Previous studies have reported that phylogenetic error is independent of incomplete taxon sampling, and that instead, longer sequences will better improve the accuracy of phylogenetic inference (Rosenberg & Kumar, 2001, 2003). In contrast, other studies have reported that fewer taxa can lead to increased variance and uncertainty in the results (Ackerly, 2000), and thus increased taxon sampling is one of the most practical ways to improve the accuracy of phylogenetic estimates (Heath et al., 2008; Hillis et al., 2003; Zwickl & Hillis, 2002). With a simulation study where random and non-random samples of taxa were drawn, it was shown that random sampling provided for statistically robust trait correlations, whereas non-random sampling (*e.g.* life-history group or investigator bias) led to a significant loss of statistical power (Ackerly, 2000). For the construction of the reduced data sets in this study, it should be noted that taxa pruning was not random, as one representative was chosen for each PV species and PV type (see Materials and Methods). Therefore, the same monophyletic lineages of PVs infecting the same hosts were subsampled for phylogenetic inference with the reduced data sets (RD1 and RD2). RD1 contains representative species with initial shorter branches as compared to those chosen for RD2. Meaning that for RD2 more diverse PV genomes are being compared, and thus RD2 represents a more greedy strategy as compared to RD1. Although it has been shown that greedy algorithms are rarely the best option for statistical power (McAuliffe et al., 2005), it has also been shown that for genomic comparisons the greedy strategy of maximizing the evolutionary divergence among species chosen from a known phylogeny can provide optimal solutions (Pardi & Goldman, 2005). Concordantly, our results indicate an effect of taxa choice in the reduced data sets, where time inference using single node calibrations with more diverse terminal taxa (RD2) leads to lower short-term evolutionary rates, than when using less diverse terminal taxa (RD1), while no differences were observed for the long-term evolutionary rates between both data sets (Fig. 2 and Table S3). When comparing the results for all data sets in this study, it also seems that RD2 renders closer estimates to those obtained for the full data sets than RD1 does (Fig. 3a-b and Tables S4-6). In the particular case of this study, increasing the number of taxa appears to reduce phylogenetic error, leading to more accurate evolutionary rate estimates, and younger basal node age estimates in the full data sets (FD and FDF). The combination of multiple data sets in our study also allows us to evaluate the impact on the estimates for deeper nodes when increasing the number of taxa in the outgroup. In this case, FD and FDF only differ in the additional presence of four fish PV genomes that were made available during the study. Indeed, the increased presence of basal fish PVs in the FDF, results in younger values for the estimate of the age of the root than any other data set (Fig. 3b). We interpret that the increased number of terminal taxa in the outgroup decreases the uncertainty about the number and nature of substitutions in this clade (Heath et al., 2008), leading to a better estimate of the age of the root.

Previous studies have shown that different substitution rates can be obtained for the same species depending on whether they are inferred on a long-term or short-term evolutionary scale. Here we demonstrate that such time-dependency of inferred evolutionary rates is also recovered for PVs. The obtained long-term evolutionary rate estimates are in line with previous overall rates obtained for the *E1* (7.1×10^-9^) and *L1* (9.6×10^-9^) genes of PVs (Shah et al., 2010). Other PV rates estimates, such as 1.84×10^-8^ for recent evolution in the HPV16 lineage (Pimenoff et al., 2017) and 1.95×10^-8^ for the evolution of a monophyletic assembly of PVs infecting felids (Rector et al., 2007), fall between the long-term and short-term rates obtained in this study. This difference can be explained by the inference of evolutionary rates in more recent times in these studies (for the HPV16 clade, using a data set spanning 0.7 Ma, and for the feline PV clade, using a data set spanning 12 Ma, respectively). Overall, the interpretation of the contrasting estimates in the literature under the framework presented here –a time-dependent behaviour for the inferred evolutionary rates for PVs– supports the hypothesis that a time dependency of the evolutionary rate is present within PVs.

With our analyses in Supplementary File 1, we show that the time-dependent rate phenomenon can be well described by fitting a power-law model to the observed data when applying individual calibration nodes. However, the increase in available statistical models and inference tools for complex evolutionary processes allowed us to incorporate the TDR model for time inference of PVs. In our analyses the TDR model performed considerably better than other models that do not allow for rate variation over time. We compared a uniformly and exponentially distributed TDR model. Even though the uniformly distributed model contained more and better dispersed transition times, we find better model fit for the exponential model. This difference in model fit may therefore imply that the rate variation through time may not be as regular as expected from a power-law function (Membrebe et al., 2019). Nevertheless, the increased complexity in the uniform TDR model (*i.e.* more transition times as compared to the exponential TDR model), can also lead to a worse fit of this model.

Regarding the evolution of substitution rates, it does not seem that PV evolution has reached an evolutionary stasis, as the PV substitution rates still vary after more than 400 million years of virus-host co-evolution (Supplementary File 1). Episodic and rare events eventually triggering an elevated substitution rate such as host jumps, switch of tissue tropism (Van Doorslaer et al., 2015; Willemsen & Bravo, 2019) and gene gain/loss events (Van Doorslaer & McBride, 2016; Willemsen et al., 2019) might prevent PVs from reaching an evolutionary stasis with their hosts, thus maintaining recent increased substitution rates.

We would like to point out that our study suffers from a number of limitations. Our phylogenetic analyses violate the assumptions of extinction (by choosing the Yule tree prior) and complete taxon sampling. In the full data sets we have only worked with reference viral genomes and have not considered the within-type viral diversity. Our jackknifing approach for the reduced data sets contributes itself to this incomplete sampling because we have explicitly chosen to not include all viral taxa. It is therefore that we tested the Yule tree prior versus the Bayesian Skyline tree prior, and the simpler Yule model rendered the best fit to the data. Consequently, the LTT plots in this study are a crude approximation to the speciation rate. Moreover, we used the host LTT plot for identifying changes in the speciation rates over time. However, the precise dating of mammalian evolution as well as the time line of speciation events are a matter of debate for specialists in the field (*e.g.* Springer et al., 2019). Therefore, our vertebrate taxa choice is biased towards species known to be the hosts of PVs for which the genome has been sequenced, and the breakpoints inferred on the host LTT plot cannot be automatically extended to match and describe the full vertebrate, amniote nor mammalian evolution. Under these limitations, the analysis of the slope time trend in the LTT plots risks of not being powerful enough and to suffer from false negative results. Notwithstanding, we were are able to identify different periods for the variation of the hots lineages through time, which are compatible with historically significant changes during the evolution of the hosts, as well as with the observed variation of viral lineages through time.

With the current set of available genomes we can distinguish four periods with different diversification rates, along the evolutionary time scale of PVs. Our results suggest that the evolutionary history of PVs is multiphasic, where subsequent radiation events allowed viruses to adapt to new niches, and a subsequent period where independent diversification between viruses and their hosts occurred within the major virus-host clades. Our results improve and refine the previous evolutionary scenario for PVs, that had suggested a biphasic evolution of PVs, where a primary radiation was directly followed by a secondary diversification event (Ignacio G. Bravo & Felez-Sanchez, 2015). The discovery of novel PVs infecting ancestral hosts, like in the present case fish, allowed for the detection of these events. Further increase in taxon sampling of both mammalian and non-mammalian hosts, together with the implementation of more flexible evolutionary models allowing for burst and decay of speciation rates, will lead to a more detailed overview of the evolutionary history of this successful viral family.

## Acknowledgements

The authors declare no conflict of interest. We are grateful to the genotoul bioinformatics platform Toulouse Midi-Pyrenees (Bioinfo Genotoul) for providing computing and storage resources. The authors acknowledge the IRD itrop HPC (South Green Platform) at IRD Montpellier for providing HPC resources that have contributed to the research results reported within this paper. This work was supported by the European Research Council Consolidator Grant CODOVIREVOL (Contract Number 647916) to IGB and by the European Union Horizon 2020 Marie Sklodowska-Curie research and innovation programme grant ONCOGENEVOL (Contract Number 750180) to AW.

## Supplementary Material

Supplementary Tables S1-S6, Supplementary Figures S1-S14, and Supplementary File 1 are available online. The alignments, R scripts, starting trees, and xml files are available at https://doi.org/10.5281/zenodo.5105951.

## Notes

https://doi.org/10.5281/zenodo.5105951

### Competing Interest Statement

The authors have declared no competing interest.

### Summary of Updates

We have uploaded additional data to Zenodo, which comes with a change of doi to the external data. This new doi has now been included in the manuscript. The figures have been labeled with figure numbers.

