## Supplementary material for "Ecological opportunity, radiation events and genomic innovations shaped the episodic evolutionary history of papillomaviruses": Fig. S1-S14

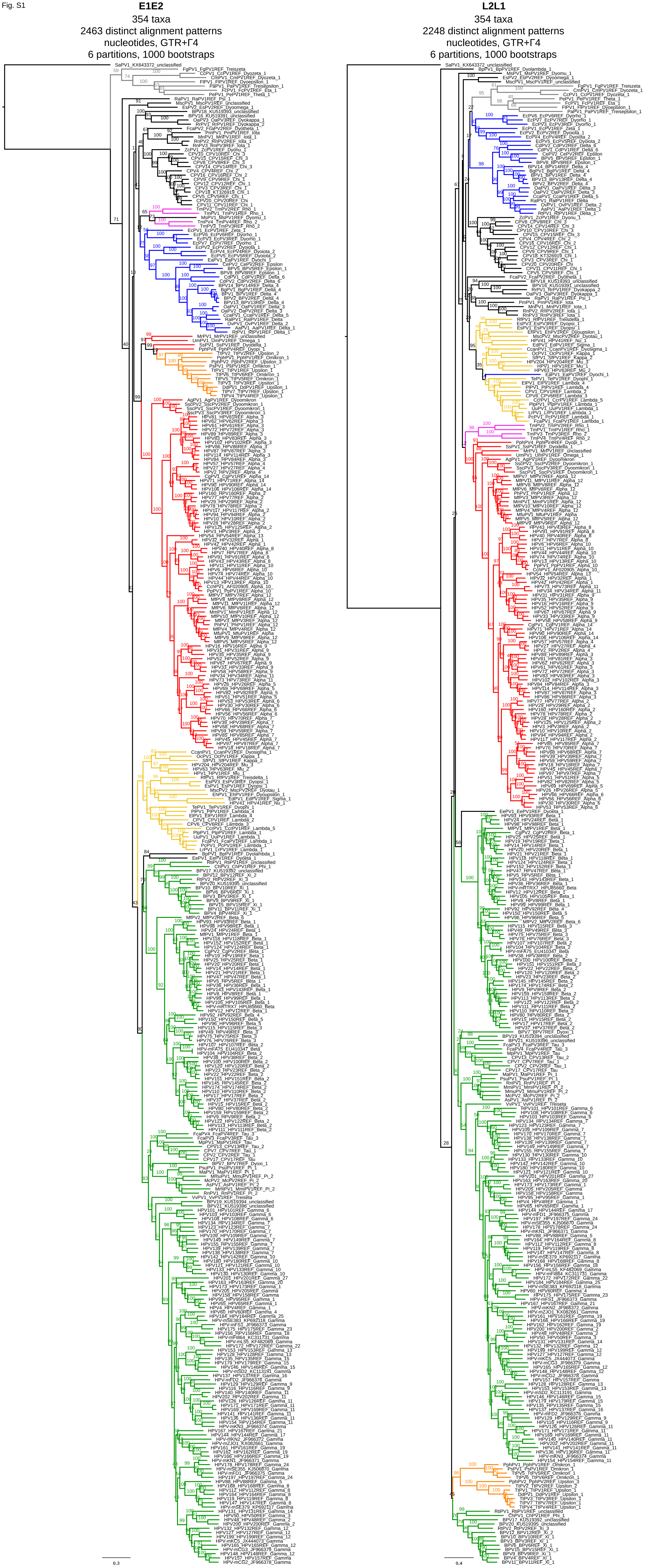


Figure S1. Maximum likelihood phylogenetic tree of the concatenated *E1-E2* and concatenated *L2-L1* nucleotide sequences of 354 PVs. Clade color codes highlight the different PV groups: red, Alpha-OmikronPVs; green, Beta-XiPVs; yellow, Lambda-MuPVs; blue, Delta-ZetaPVs; gray, a yet unclassified crown group consisting of PVs infecting birds and turtles; purple, a yet unclassified clade consisting of PVs infecting manatees; and orange, recombinant PVs infecting Cetaceans. These recombinant PVs infecting appear to have an early gene region that resembles a non-recombinant cetacean PV in the Alpha-Omikron crown group (colored red) and a late gene region that resembles XiPVs infecting Bovidae and Cervidae in the Beta-Xi crown group (colored green). Values on branches correspond to ML bootstrap support values.

Fig. S2

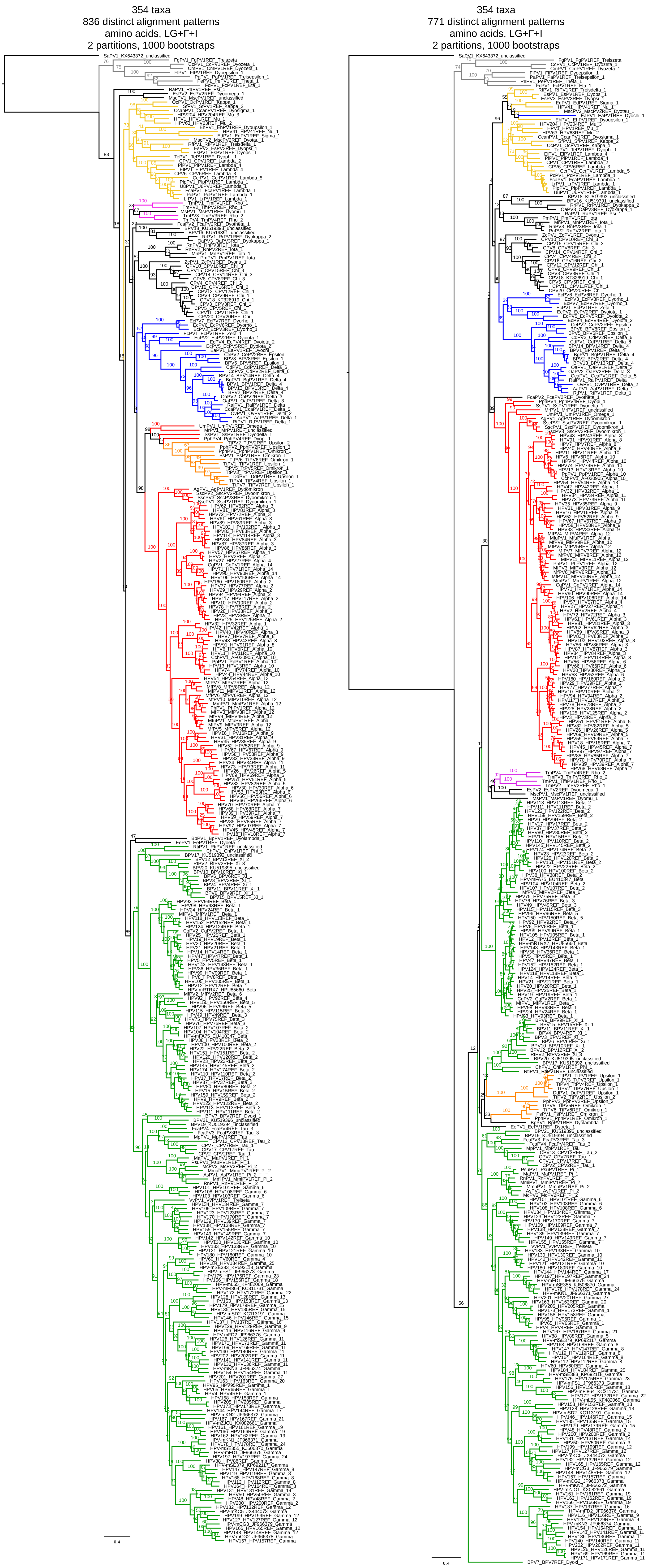


Figure S2. Maximum likelihood phylogenetic tree of the concatenated *E1-E2* and concatenated *L2-L1* amino acid sequences of 354 PVs. Clade color codes highlight the different PV groups: red, Alpha-OmikronPVs; green, Beta-XiPVs; yellow, Lambda-MuPVs; blue, Delta-ZetaPVs; gray, a yet unclassified crown group consisting of PVs infecting birds and turtles; purple, a yet unclassified clade consisting of PVs infecting manatees; and orange, recombinant PVs infecting Cetaceans. These recombinant PVs infecting appear to have an early gene region that resembles a non-recombinant cetacean PV in the Alpha-Omikron crown group (colored red) and a late gene region that resembles XiPVs infecting Bovidae and Cervidae in the Beta-Xi crown group (colored green). Values on branches correspond to ML bootstrap support values.

Fig. S3

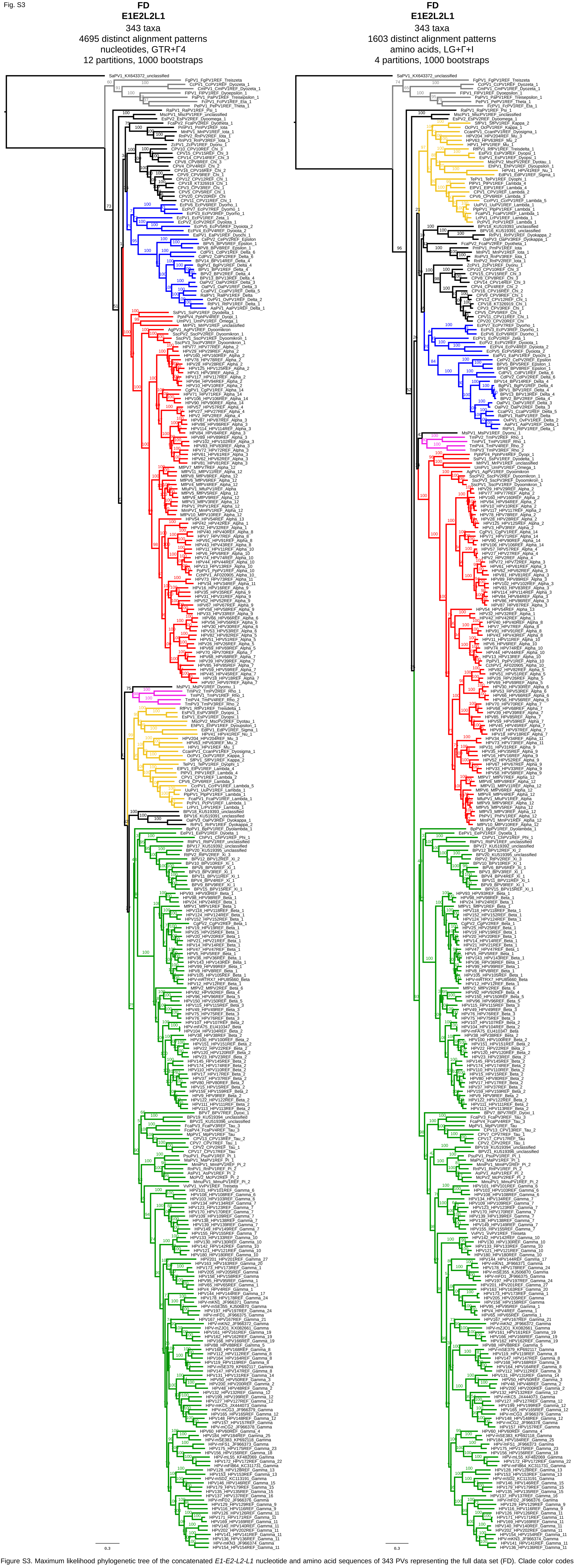


Figure S3. Maximum likelihood phylogenetic tree of the concatenated *E1-E2-L2-L1* nucleotide and amino acid sequences of 343 PVs representing the full data set (FD). Clade color codes highlight the different PV groups: red, Alpha-OmikronPVs; green, Beta-XiPVs; yellow, Lambda-MuPVs; blue, Delta-ZetaPVs; gray, a yet unclassified crown group consisting of PVs infecting birds and turtles; and purple, a yet unclassified clade consisting of PVs infecting manatees. Values on branches correspond to ML bootstrap support values.

Fig. S4

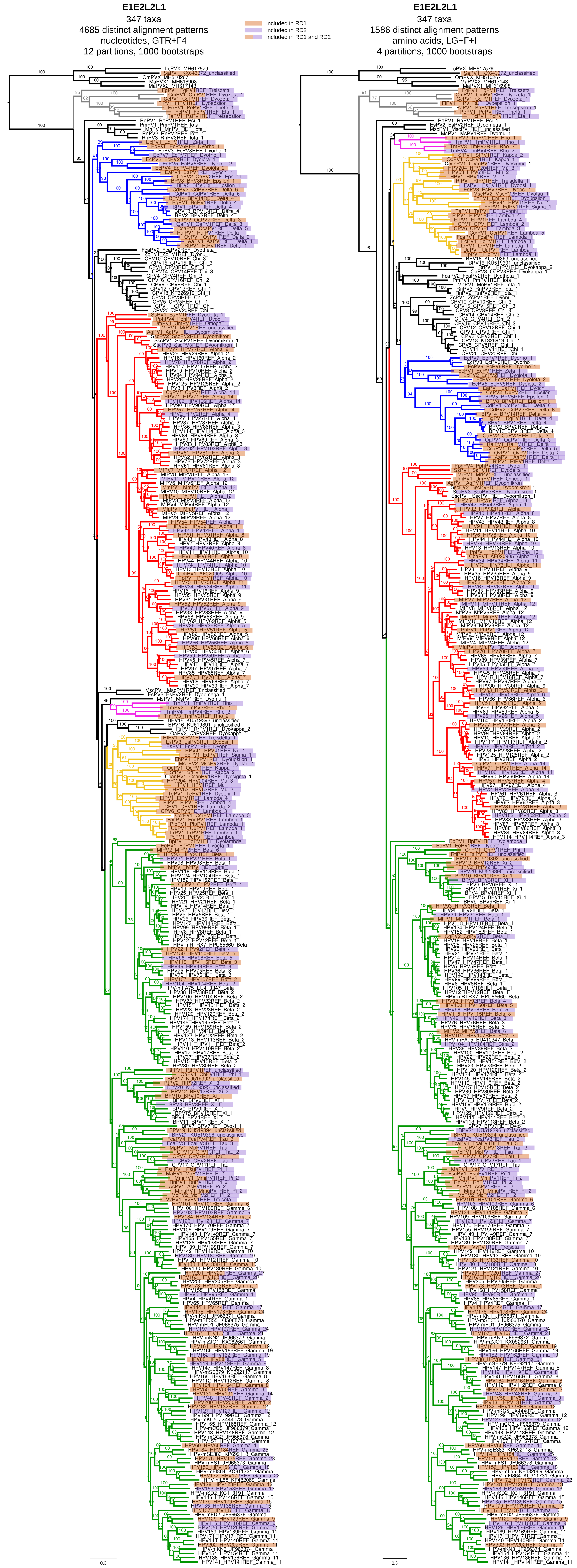


Figure S4. Maximum likelihood phylogenetic tree of the concatenated *E1-E2-L2-L1* nucleotide and amino acid sequences of 347 PVs representing the full data set plus later published fish PV genomes (FDF). Clade color codes highlight the different PV groups: red, Alpha-OmikronPVs; green, Beta-XiPVs; yellow, Lambda-MuPVs; blue, Delta-ZetaPVs; gray, a yet unclassified crown group consisting of PVs infecting birds and turtles; and purple, a yet unclassified clade consisting of PVs infecting manatees. Values on branches correspond to ML bootstrap support values. The taxa included in RD1 (Fig. S5) and/or RD2 (Fig. S6) are highlighted according to the color code indicated in the legend.

Fig. S5

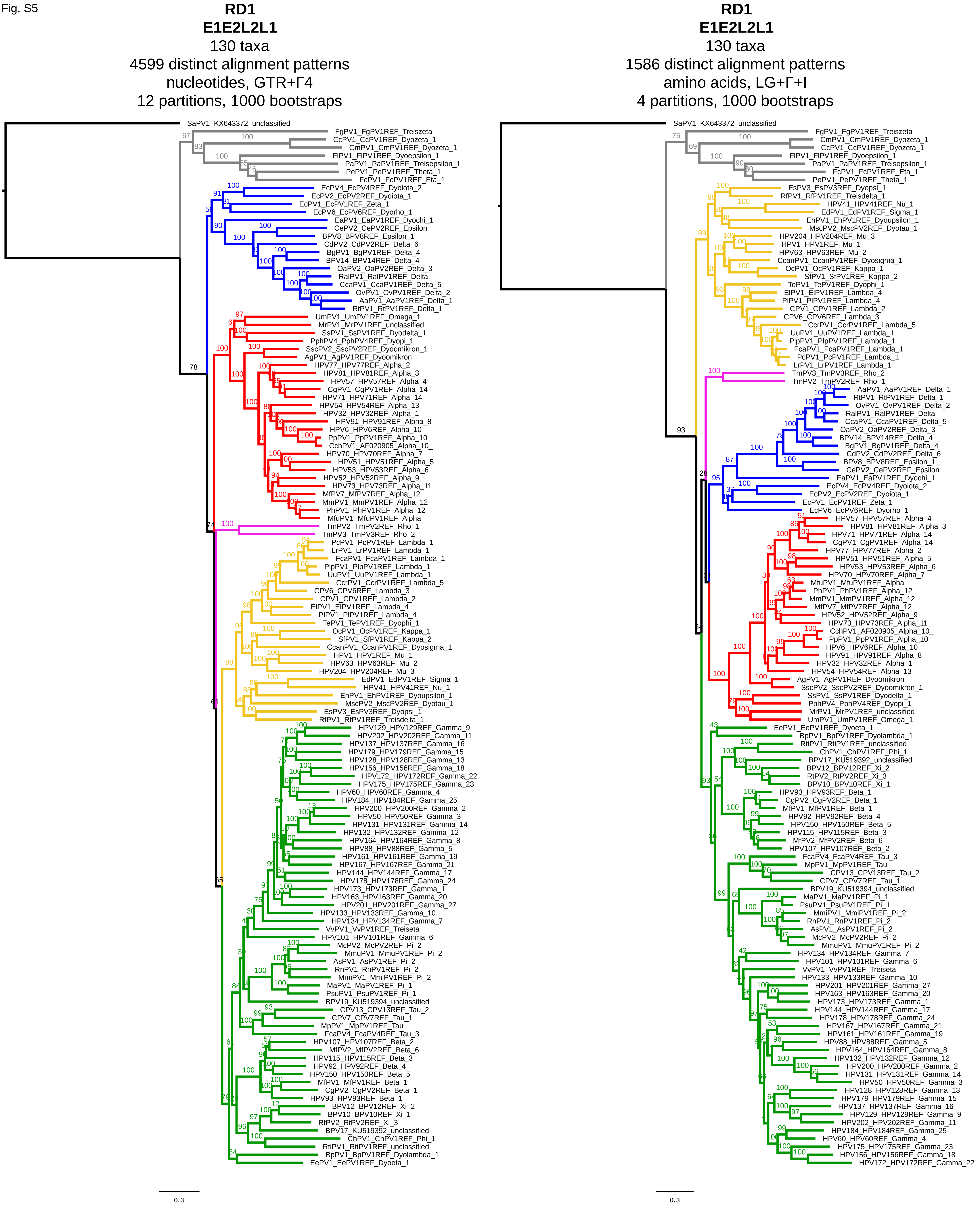


Figure S5. Maximum likelihood phylogenetic tree of the concatenated *E1-E2-L2-L1* nucleotide and amino acid sequences of 130 PVs in representative data set 1 (RD1). Clade color codes highlight the different PV groups: red, Alpha-OmikronPVs; green, Beta-XiPVs; yellow, Lambda-MuPVs; blue, Delta-ZetaPVs; gray, a yet unclassified crown group consisting of PVs infecting birds and turtles; and purple, a yet unclassified clade consisting of PVs infecting manatees. Values on branches correspond to ML bootstrap support values.

Fig. S6

**RD2**
**E1E2L2L1**
130 taxa
4597 distinct alignment patterns
nucleotides, GTR+Γ4
12 partitions, 1000 bootstraps

**RD2**
**E1E2L2L1**
130 taxa
1585 distinct alignment patterns
amino acids, LG+Γ+I
4 partitions, 1000 bootstraps

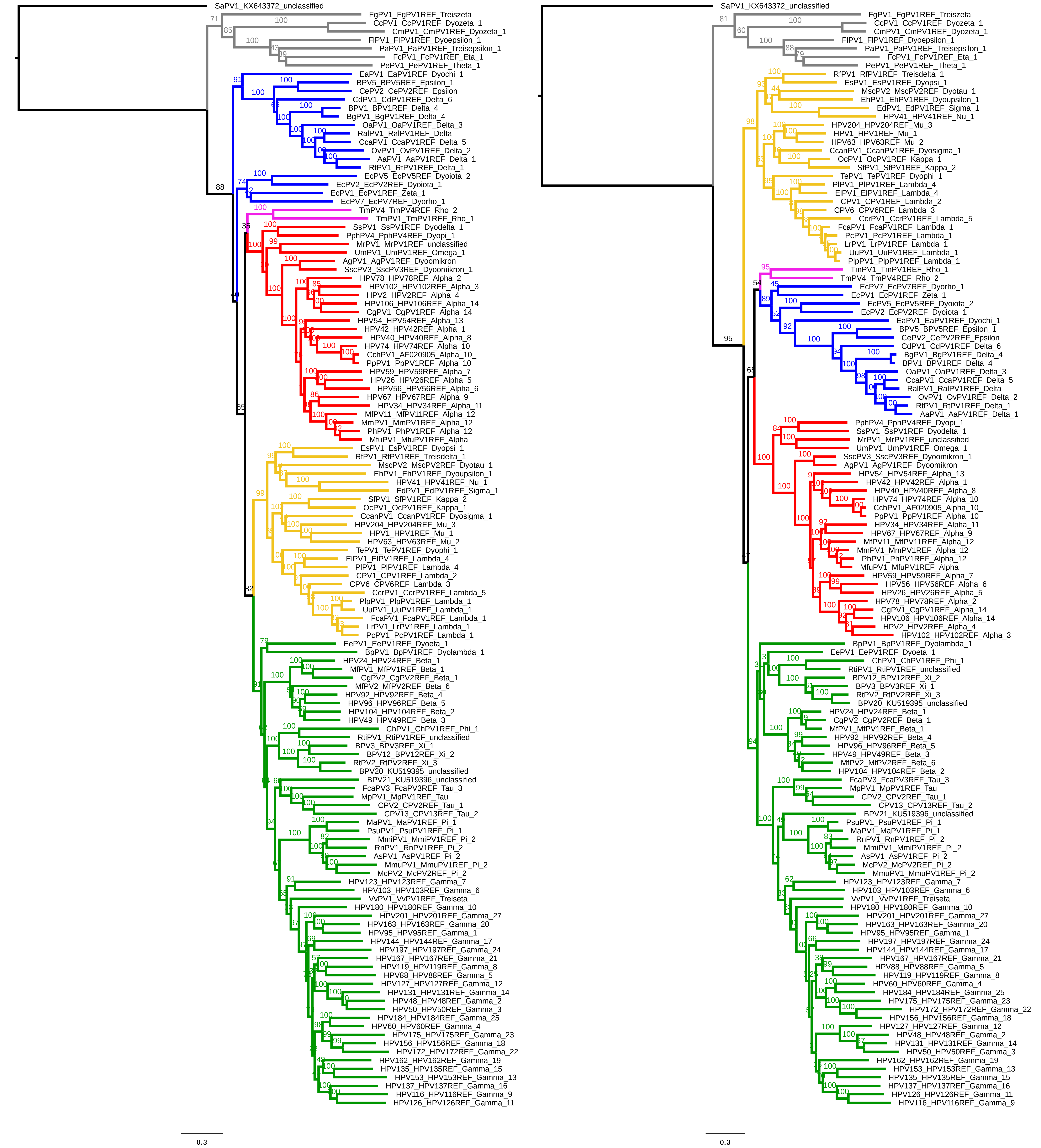


Figure S6. Maximum likelihood phylogenetic tree of the concatenated *E1-E2-L2-L1* nucleotide and amino acid sequences of 130 PVs in representative data set 2 (RD2). Clade color codes highlight the different PV groups: red, Alpha-OmikronPVs; green, Beta-XiPVs; yellow, Lambda-MuPVs; blue, Delta-ZetaPVs; gray, a yet unclassified crown group consisting of PVs infecting birds and turtles; and purple, a yet unclassified clade consisting of PVs infecting manatees. Values on branches correspond to ML bootstrap support values.

Fig. S7

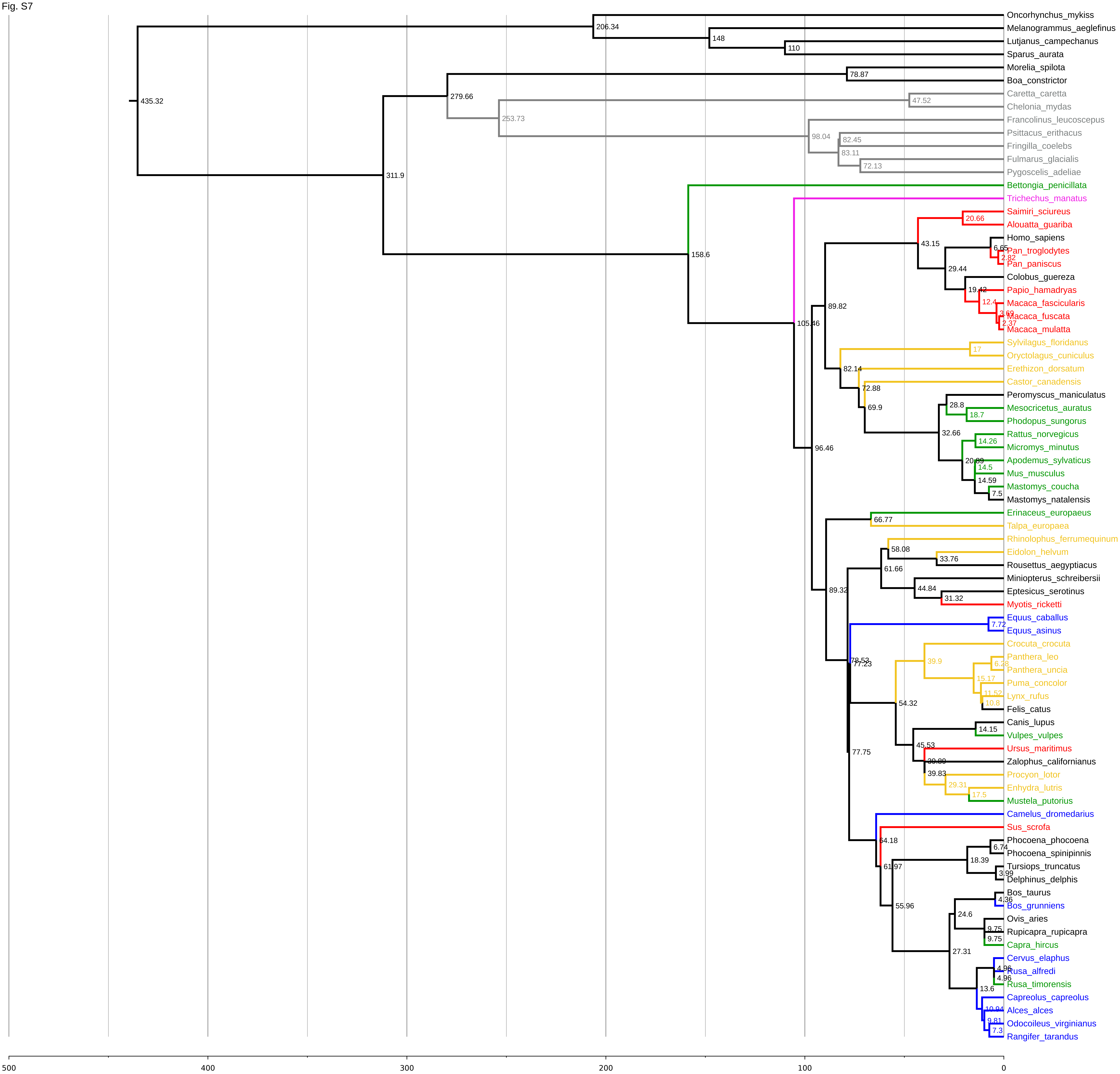


Figure S7. Dated phylogenetic tree for the known host species of PVs. The tree was constructed was constructed using TimeTree (http://www.timetree.org/), by providing a list of all PV host species included in this study. The scale bar is given in million years ago (Ma). Values at the nodes correspond to the diverge time estimate in Ma. Branches leading leading to taxa and taxa names that are unique to a specific PV clade are colored according to the PV crown-group classification: red, Alpha-OmikronPVs; green, Beta-XiPVs; yellow, Lambda-MuPVs; blue, Delta-ZetaPVs; gray, a yet unclassified crown group consisting of PVs infecting birds and turtles; and purple, a yet unclassified clade consisting of PVs infecting manatees.

Fig. S8

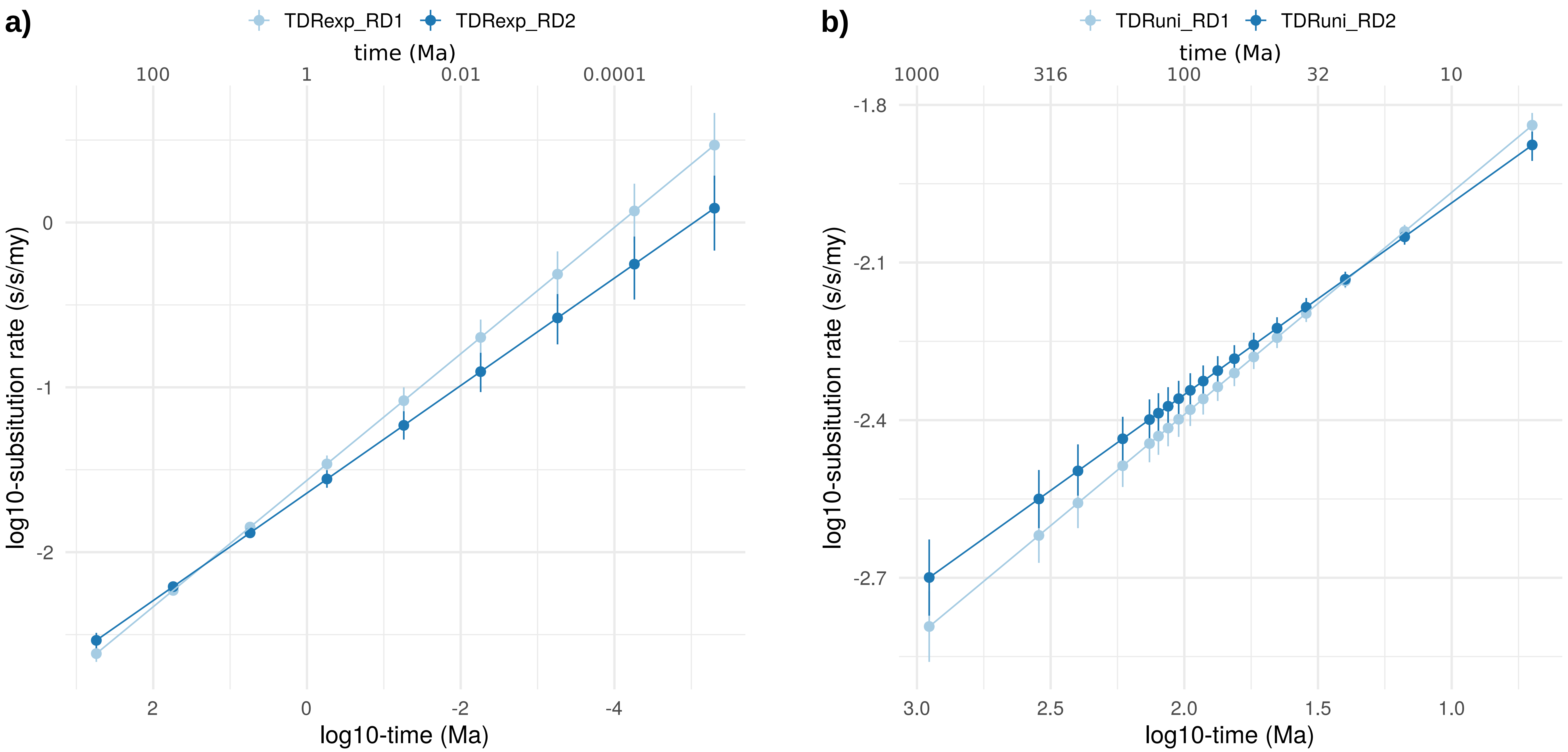


Figure S8. Time-dependent rate effect under both an epoch structure with exponentially (a) and uniformly (b) distributed time intervals. The x-axis represents time from past to present in million years ago (Ma). The y-axis represents the evolutionary rate in substitutions/site/million years (s/s/my). Both axes are on a log10 scale. The error bars represent the 95% highest posterior density (HPD) intervals of the rate estimates.

Fig. S9

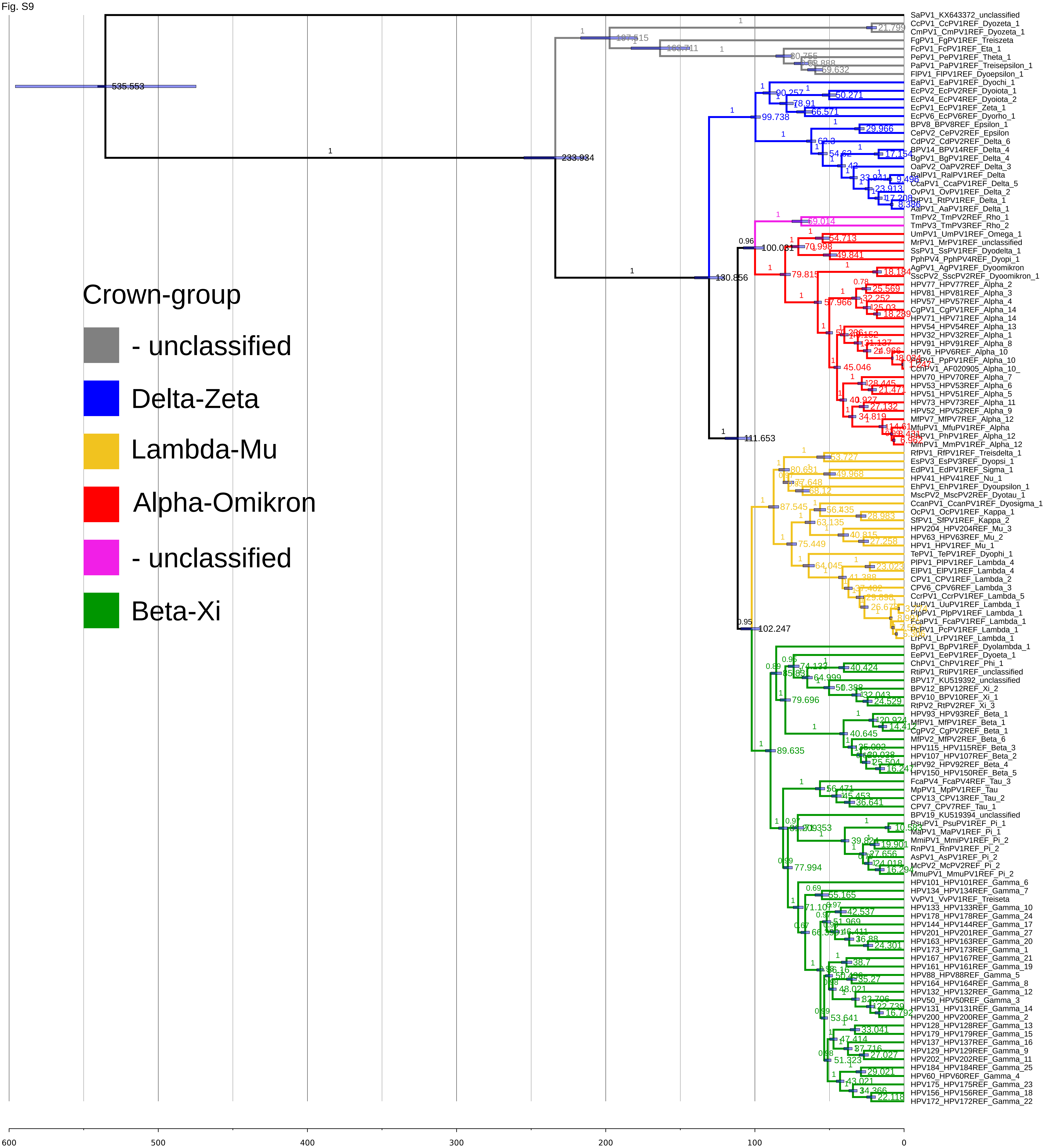


Figure S9. Dated Bayesian phylogenetic tree for the reduced representative data set 1 (RD1) containing 130 PVs. The tree was constructed at the nucleotide level based on the concatenated *E1-E2-L2-L1* genes. The scale bar is given in million years ago (Ma). Values at the nodes correspond to the age and posterior probabilities are indicated on the branches. Error bars encompass 95% HPD for the age of the nodes. The clades are colored according to the PV crown-group classification, as indicated in the legend.

Fig. S10

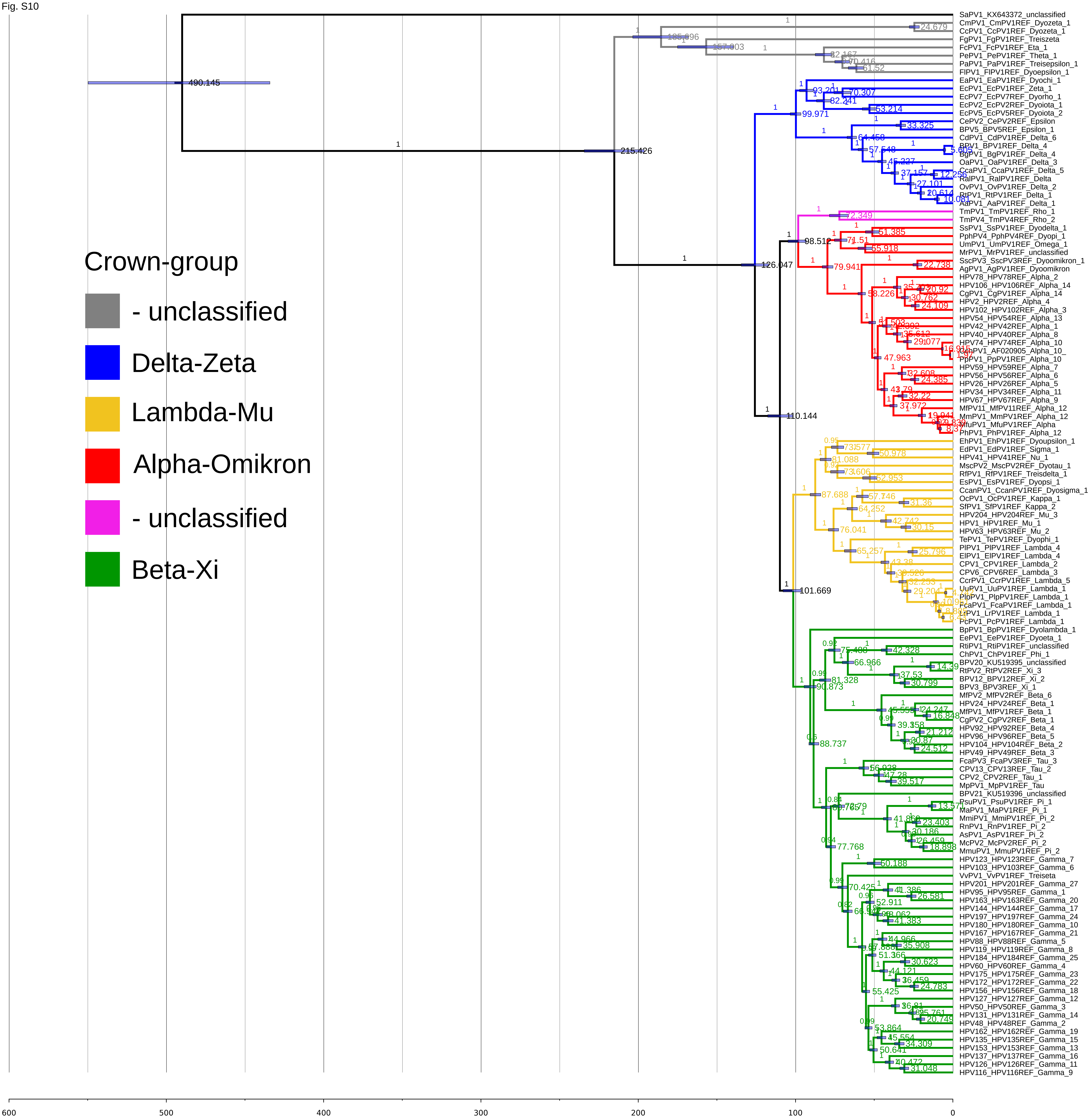


Figure S10. Dated Bayesian phylogenetic tree for the reduced representative data set 2 (RD2) containing 130 PVs. The tree was constructed at the nucleotide level based on the concatenated *E1-E2-L2-L1* genes. The scale bar is given in million years ago (Ma). Values at the nodes correspond to the age and posterior probabilities are indicated on the branches. Error bars encompass 95% HPD for the age of the nodes. The clades are colored according to the PV crown-group classification, as indicated in the legend.

Fig. S11

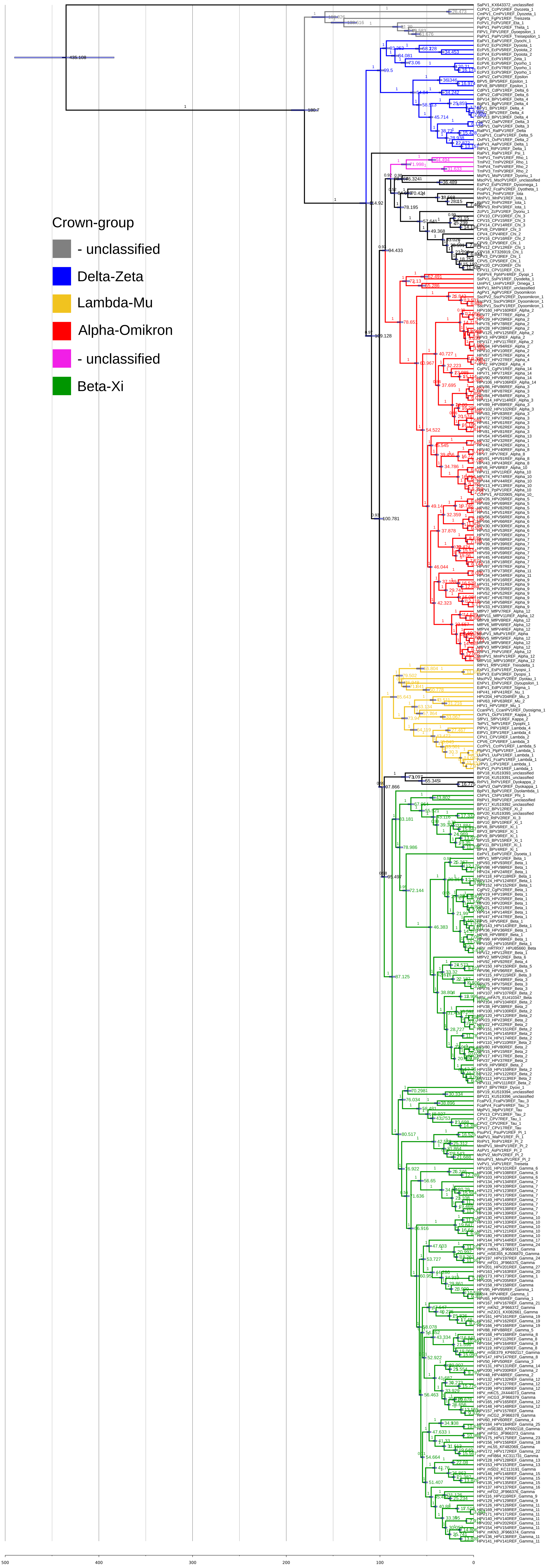


Figure S11. Dated Bayesian phylogenetic tree for the full data set (FD) containing 343 PVs. The tree was constructed at the nucleotide level based on the concatenated *E1-E2-L2-L1* genes. The scale bar is given in million years ago (Ma). Values at the nodes correspond to the age and posterior probabilities are indicated on the branches. Error bars encompass 95% HPD for the age of the nodes. The clades are colored according to the PV crown-group classification, as indicated in the legend.

Fig. S12

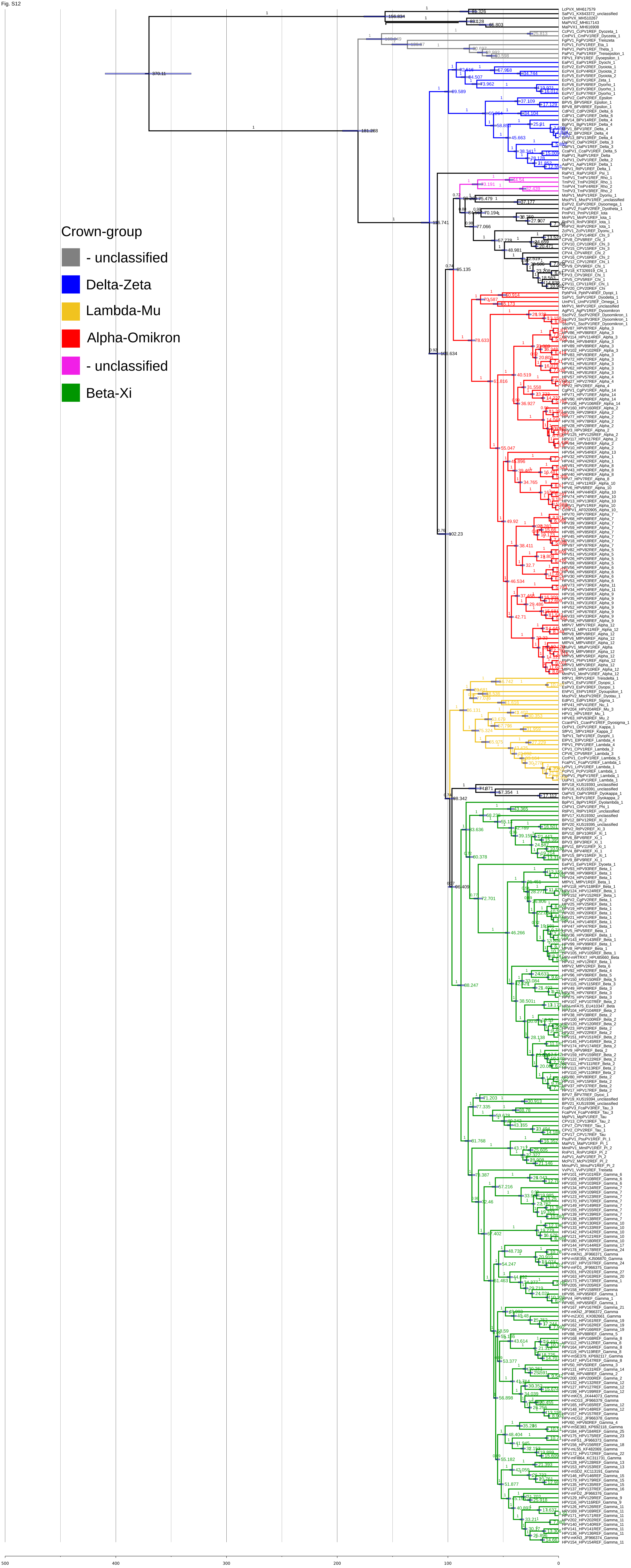


Figure S12. Dated Bayesian phylogenetic tree for the full data set (FDF) containing 347 PVs. The tree was constructed at the nucleotide level based on the concatenated *E1-E2-L2-L1* genes. The scale bar is given in million years ago (Ma). Values at the nodes correspond to the age and posterior probabilities are indicated on the branches. Error bars encompass 95% HPD for the age of the nodes. The clades are colored according to the PV crown-group classification, as indicated in the legend.

Fig. S13

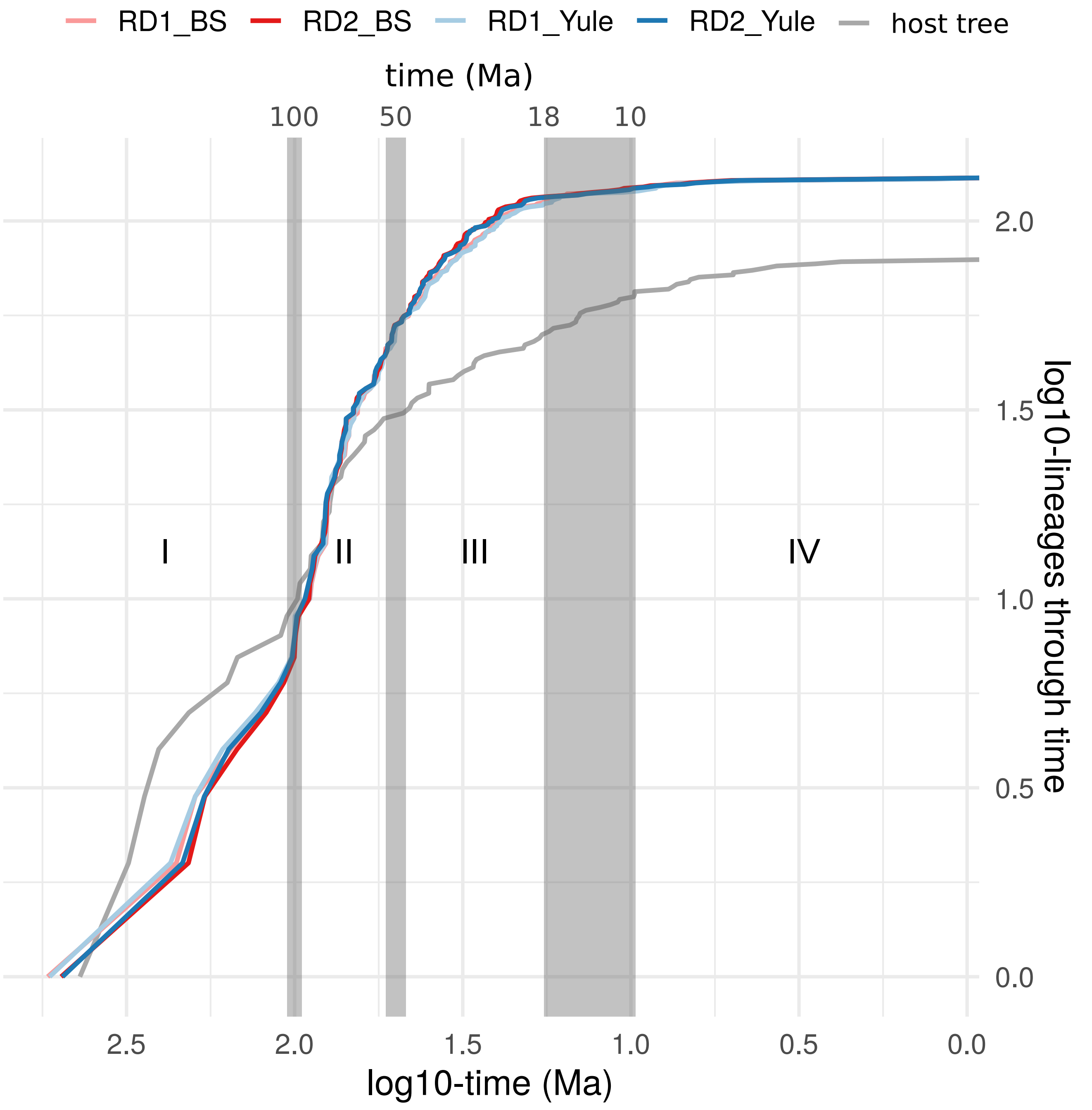


Figure S13. Lineages through time (LTT) plot for RD1, RD2 and the host tree. The x-axis represents time from past to present in Ma. The y-axis represents the number of lineages. Both axes are on a log10 scale. The trees were constructed using a Bayesian Skyline (BS) or a Yule tree prior. The dotted lines indicate separate periods (I-IV). This figure demonstrates that both tree priors give highly similar results. .

Fig. S14

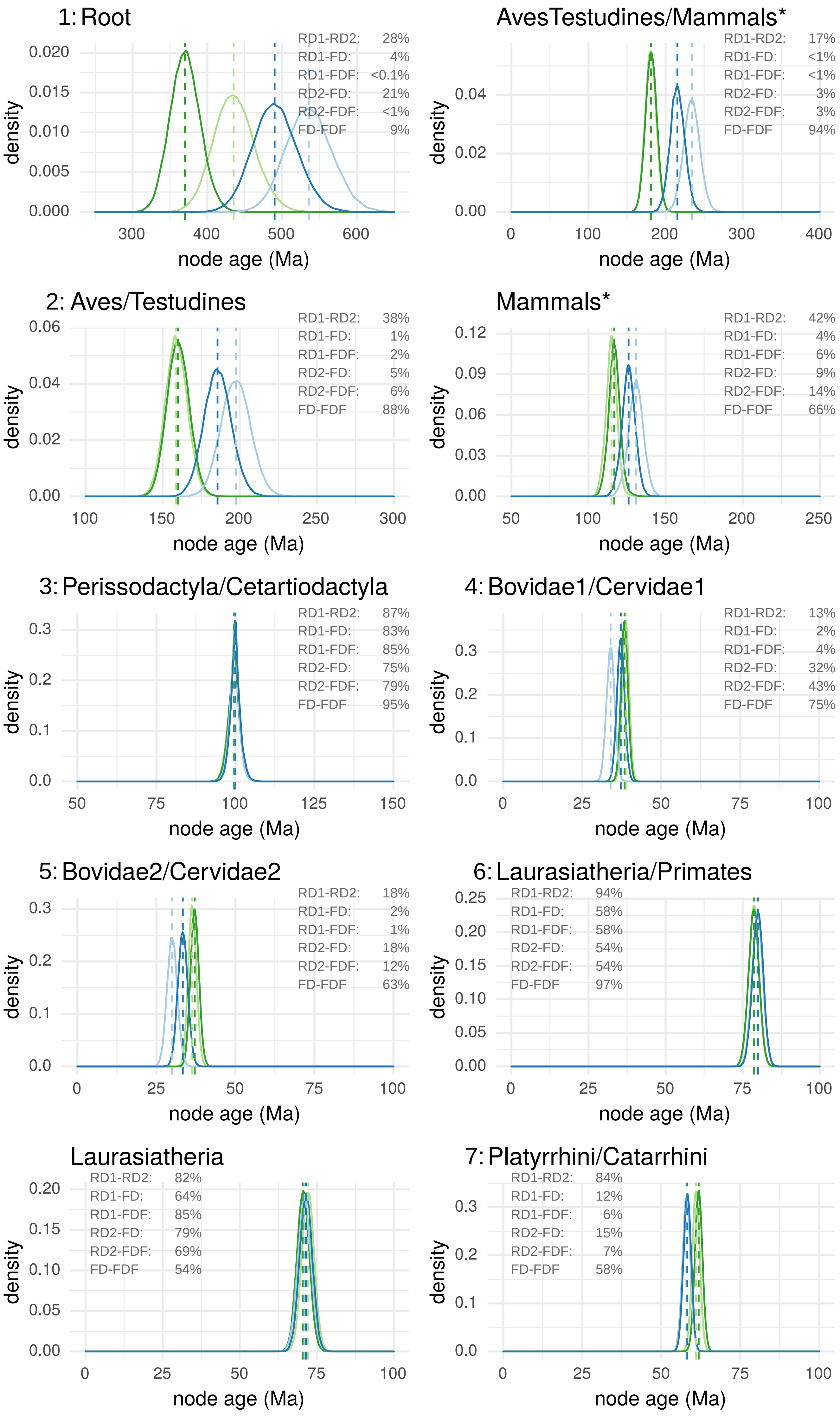
1: Root
RD1-RD2: 28%
RD1-FD: 4%
RD1-FDF: <0.1%
RD2-FD: 21%
RD2-FDF: <1%
FD-FDF 9%
density
node age (Ma)
AvesTestudines/Mammals*
RD1-RD2: 17%
RD1-FD: <1%
RD1-FDF: <1%
RD2-FD: 3%
RD2-FDF: 3%
FD-FDF 94%
density
node age (Ma)
2: Aves/Testudines
RD1-RD2: 38%
RD1-FD: 1%
RD1-FDF: 2%
RD2-FD: 5%
RD2-FDF: 6%
FD-FDF 88%
density
node age (Ma)
Mammals*
RD1-RD2: 42%
RD1-FD: 4%
RD1-FDF: 6%
RD2-FD: 9%
RD2-FDF: 14%
FD-FDF 66%
density
node age (Ma)
3: Perissodactyla/Cetartiodactyla
RD1-RD2: 87%
RD1-FD: 83%
RD1-FDF: 85%
RD2-FD: 75%
RD2-FDF: 79%
FD-FDF 95%
density
node age (Ma)
4: Bovidae1/Cervidae1
RD1-RD2: 13%
RD1-FD: 2%
RD1-FDF: 4%
RD2-FD: 32%
RD2-FDF: 43%
FD-FDF 75%
density
node age (Ma)
5: Bovidae2/Cervidae2
RD1-RD2: 18%
RD1-FD: 2%
RD1-FDF: 1%
RD2-FD: 18%
RD2-FDF: 12%
FD-FDF 63%
density
node age (Ma)
6: Laurasiatheria/Primates
RD1-RD2: 94%
RD1-FD: 58%
RD1-FDF: 58%
RD2-FD: 54%
RD2-FDF: 54%
FD-FDF 97%
density
node age (Ma)
Laurasiatheria
RD1-RD2: 82%
RD1-FD: 64%
RD1-FDF: 85%
RD2-FD: 79%
RD2-FDF: 69%
FD-FDF 54%
density
node age (Ma)
7: Platyrrhini/Catarrhini
RD1-RD2: 84%
RD1-FD: 12%
RD1-FDF: 6%
RD2-FD: 15%
RD2-FDF: 7%
FD-FDF 58%
density
node age (Ma)

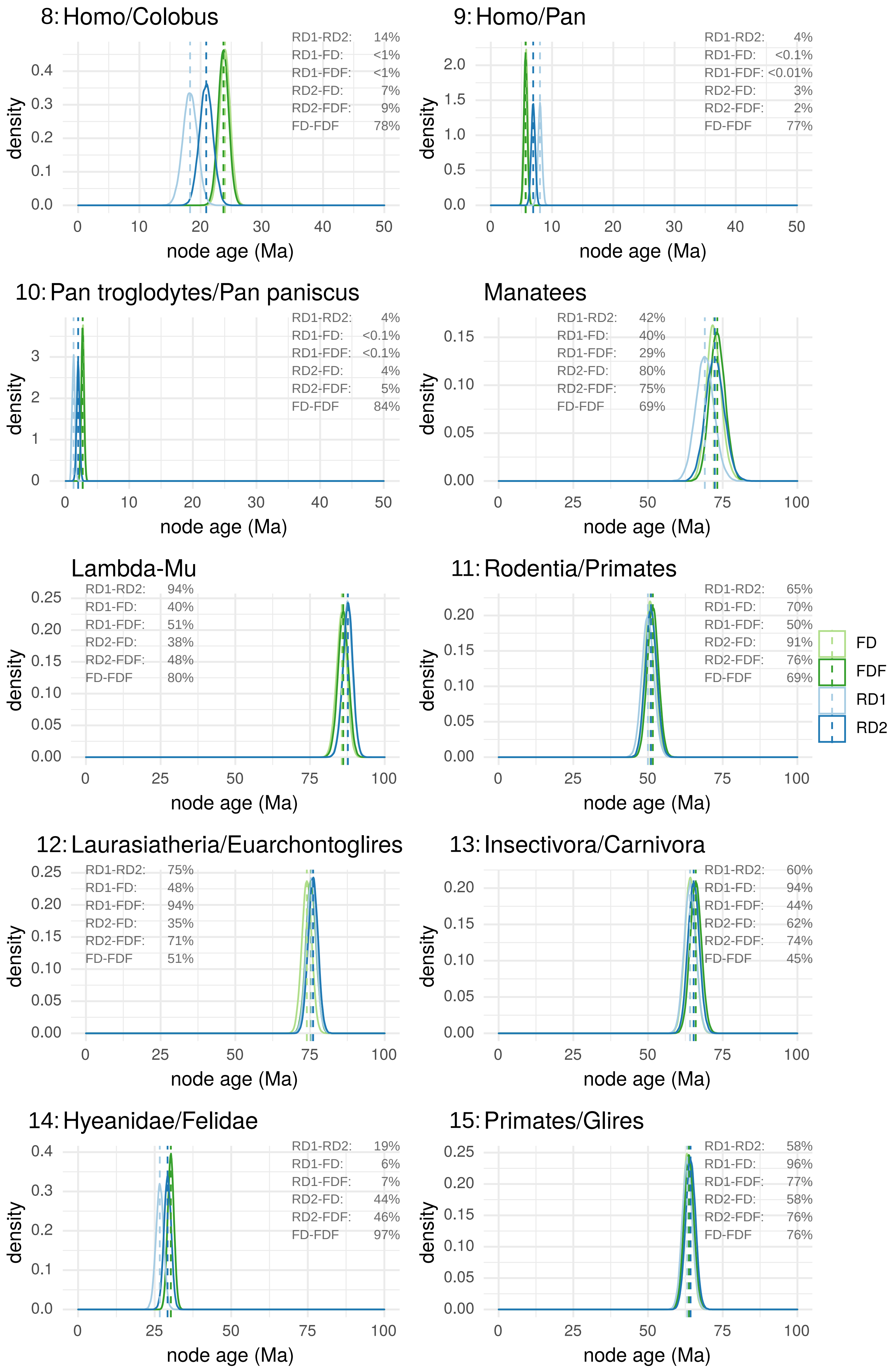
8: Homo/Colobus
RD1-RD2: 14%
RD1-FD: <1%
RD1-FDF: <1%
RD2-FD: 7%
RD2-FDF: 9%
FD-FDF 78%
9: Homo/Pan
RD1-RD2: 4%
RD1-FD: <0.1%
RD1-FDF: <0.01%
RD2-FD: 3%
RD2-FDF: 2%
FD-FDF 77%
10: Pan troglodytes/Pan paniscus
RD1-RD2: 4%
RD1-FD: <0.1%
RD1-FDF: <0.1%
RD2-FD: 4%
RD2-FDF: 5%
FD-FDF 84%
Manatees
RD1-RD2: 42%
RD1-FD: 40%
RD1-FDF: 29%
RD2-FD: 80%
RD2-FDF: 75%
FD-FDF 69%
Lambda-Mu
RD1-RD2: 94%
RD1-FD: 40%
RD1-FDF: 51%
RD2-FD: 38%
RD2-FDF: 48%
FD-FDF 80%
11: Rodentia/Primates
RD1-RD2: 65%
RD1-FD: 70%
RD1-FDF: 50%
RD2-FD: 91%
RD2-FDF: 76%
FD-FDF 69%
12: Laurasiatheria/Euarchontoglires
RD1-RD2: 75%
RD1-FD: 48%
RD1-FDF: 94%
RD2-FD: 35%
RD2-FDF: 71%
FD-FDF 51%
13: Insectivora/Carnivora
RD1-RD2: 60%
RD1-FD: 94%
RD1-FDF: 44%
RD2-FD: 62%
RD2-FDF: 74%
FD-FDF 45%
14: Hyeanidae/Felidae
RD1-RD2: 19%
RD1-FD: 6%
RD1-FDF: 7%
RD2-FD: 44%
RD2-FDF: 46%
FD-FDF 97%
15: Primates/Glires
RD1-RD2: 58%
RD1-FD: 96%
RD1-FDF: 77%
RD2-FD: 58%
RD2-FDF: 76%
FD-FDF 76%
density
node age (Ma)
FD
FDF
RD1
RD2

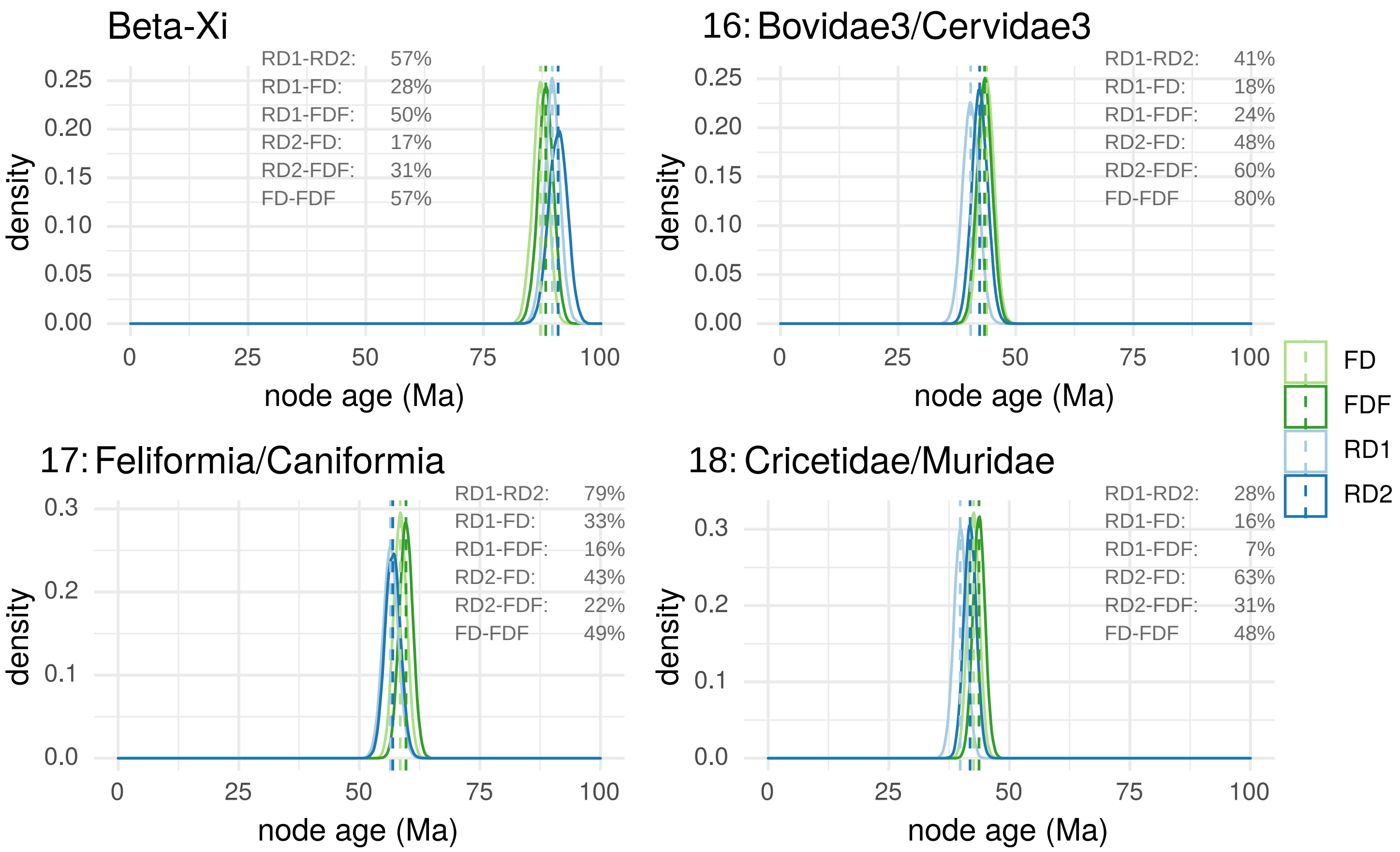


Figure S14. Comparison of the posterior distributions of tmrca estimates for different clades along the PV tree. Time inference was performed with an exponential TDR model and a Yule tree prior, using the RD1, RD2, FD and FDF data sets. The x-axis represents the node age (Ma), for which the names are indicated above each plot. The numbers next to the node names, indicate the nodes used for calibration. The y-axis represents the density of the data. The median of each density distribution is indicated with a vertical dotted line. The percentage scores indicate to which extent the density distributions overlap.
