## Supplementary File 1 for "Ecological opportunity, radiation events and genomic innovations shaped the episodic evolutionary history of papillomaviruses"

In this document we describe how we verified that a time-dependency of molecular rate estimates exists for PVs, before the implementation of a time-dependent rate clock model in BEAST (Membrebe et al., 2019). For the results shown in this supplementary file, bayesian time inference was performed at the nucleotide level using BEAST v.1.8.3 (Drummond et al., 2012), under the GTR+Γ4 model, using twelve partitions, the uncorrelated relaxed clock model (Drummond et al., 2006) with a lognormal distribution and a continuous quantile parametrisation (Li & Drummond, 2012), and the Yule speciation process tree prior (Gernhard, 2008; Yule, 1925). For each calibration node, a normal distribution was assumed with the host divergence estimate and standard deviation based on the confidence intervals indicated in Table 1 in main text.

First we performed time inference on the different data sets (RD1, RD2, FD and FDF) using all 18 calibration points together. For all four data sets, we obtained an overall substitution rate of around $8x10^{-9}$ substitutions per site per year (Table SF1).

| | **Data set** | **Mean** | **Median** | **95% HPD** | |
|---|---|---|---|---|---|
| **multiple calibrations** | RD1 | $7.90x10^{-9}$ | $7.89x10^{-9}$ | ($7.45x10^{-9}$; | $8.35x10^{-9}$) |
| | RD2 | $8.01x10^{-9}$ | $8.01x10^{-9}$ | ($7.55x10^{-9}$; | $8.47x10^{-9}$) |
| | FD | $7.99x10^{-9}$ | $7.98x10^{-9}$ | ($7.62x10^{-9}$ ; | $8.36x10^{-9}$) |
| | FDF | $7.68x10^{-9}$ | $7.67x10^{-9}$ | ($7.31x10^{-9}$; | $8.02x10^{-9}$) |
| **single calibrations** | RD1 | $9.56x10^{-9}$ | $7.39x10^{-9}$ | NA | |
| | RD2 | $6.93x10^{-9}$ | $6.71x10^{-9}$ | NA | |

*Table SF1. Overall substitution rates of evolution for PVs obtained by using different data sets. The substitution rates were inferred by (i) constructing one dated bayesian phylogenetic tree for each data set (RD1, RD2, FD, and FDF) using all 18 calibrations points together (multiple calibrations), and by (ii) constructing 18 dated bayesian phylogenetic trees for RD1 and RD2 using each calibration point separately (single calibrations). For the single calibrations the mean and median values were calculated on the 18 combined posterior distributions. Values are in substitutions per site per year.*

For each of the eight sub-clades of the dated FDF tree, the mean age between the internal nodes was calculated and displayed with a density graph (Fig. SF1a). The densities were compared with a permutation test of equality using 100 bootstraps as well as with a Kruskal-Wallis rank sum test. Significant differences were found between the clades ($chi^2$ = 60.202, *df* = 7, *p* = 1.376e-10), where the distribution of the *AlphaPVs* is significantly different from that of the *GammaPVs* ($chi^2$ = 27.552, *df* = 1, *p* = 1.529e-07), and all PV genera are different from the basal *Aves-Testudines* and *Fish* PV clades. Subsequently, the substitution rate was obtained for the internal nodes of the dated FDF tree (Fig. SF1b). As a general trend, we observe that younger nodes have a higher substitution rate as compared to older nodes (Pearson's product-moment correlation FDF: -0.157). This relationship is significant when fitted to a linear model, however this model does not explain well the data (adj. $R^2$= 0.022, *F* = 8.668, *df* = 343, *p* = 0.003). When stratifying the data in the eight PV sub-clades, we observe that these PV groups have striking different

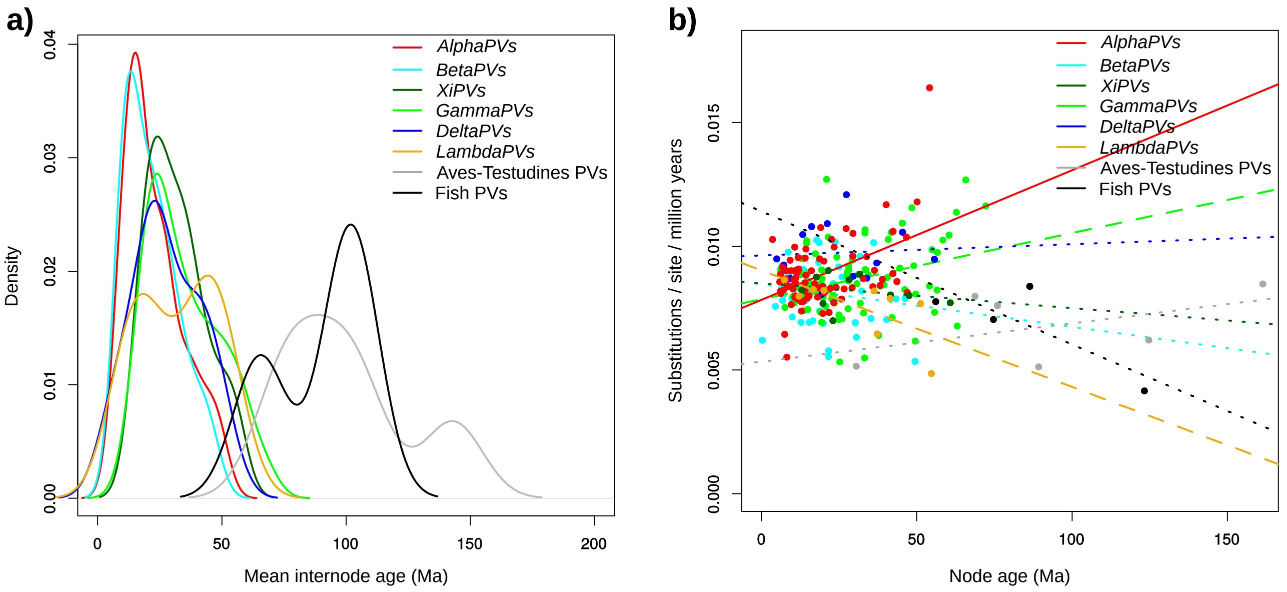


*Figure SF1. Internode age and substitution rate for eight PV clades in the FDF dataset. The PV clades are indicated in the legend, where PV genera are written in italics and unclassified clades are named according to the corresponding host species.* ***a)*** *Density graph of the mean age between internal nodes of the PV clades.* ***b)*** *Relationship of the node age and the substitution rate of the PV clades. A straight line was fitted to the data point for each of the clades. The continuous line indicates a significant trend (AlphaPVs: adj. $R^2$ = 0.215, F = 22.070, df = 76, p = 1.147e-05), dashed lines indicate a near significant trend (XiPVs and GammaPVs), and dotted lines indicate no significant trend.*

trends, where the *Alpha*- and *GammaPVs* actually display lower substitution rates in younger nodes as compared to older nodes. The highest substitution rate is present within *AlphaPVs* (Fig. SF1b). This sudden increase in substitution rate coincides with the appearance of the *E5* oncogene within the *AlphaPV* clade (Willemsen & Bravo, 2019). Although this high rate might contribute to the significant linear relationship between the node age and substitution rate within *AlphaPVs*, this relationship remains significant upon eliminating this point (adj. $R^2 = 0.124$, $F = 11.74$, $df = 75$, $p < 0.001$).

To verify the effect of the node age on the molecular rate estimates, phylogenetic inference was performed separately with each calibration point on both RD1 and RD2. Thus, per data set, 18 independent time inferences were performed. As shown in Fig. SF2a and SF2b, there is a negative correlation (RD1: *Spearman's rho* = -0.8225, *S* = 1766, *p* = 2.419e-05; RD2: *Spearman's rho* = -0.5170, *S* = 1470, *p* = 0.0299) between the substitution rate (substitutions/site/million years) and the inferred time. The data set used also has an effect on the substitution rate, where in RD1 the rates range from $4.2 \times 10^{-9}$ to $2.4 \times 10^{-8}$ substitutions per site per year, while in RD2 the upper value of this range is more than two times lower, ranging from $4.5 \times 10^{-9}$ to $1.0 \times 10^{-8}$. Nonetheless, when using single calibration points, the median substitution rates are similar for RD1 and RD2, and approximate those obtained when using all calibration points together (Table SF1). Altogether, these results show that younger nodes tend to render higher substitution rates compared to older nodes, a typical signature of a time-dependent rate phenomenon (TDRP) (Aiewsakun & Katzourakis, 2015; Duchene et al., 2014; Ho et al., 2007).

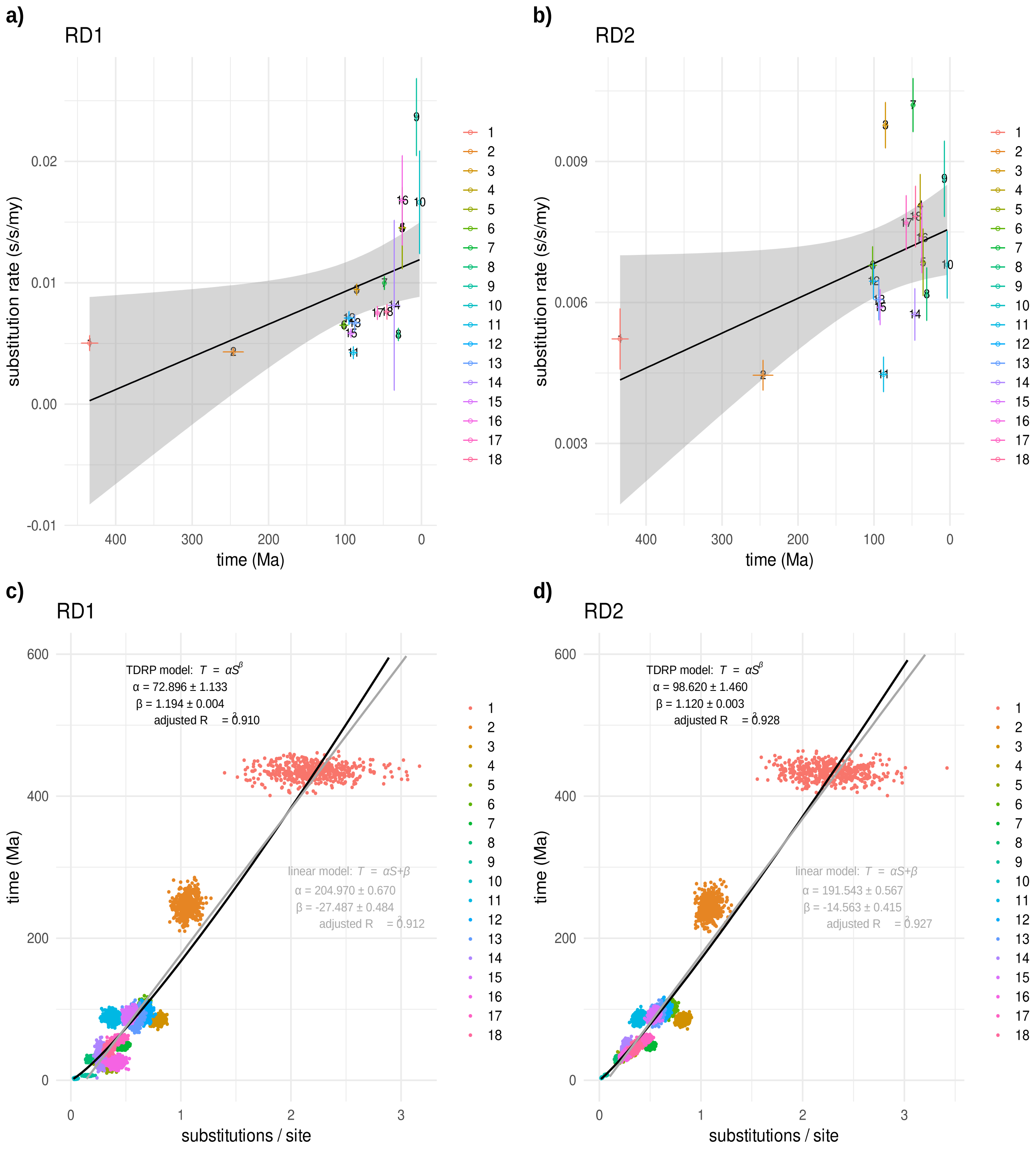


*Figure SF2. Time-dependent rate phenomenon for PVs.* ***a)*** *Substitution rates and time were inferred independently on 18 calibration points using RD1. Therefore, 18 dated bayesian phylogenetic trees were constructed, each with a single calibration point. For each calibration point, the mean and sd of the inferred time in million years ago (Ma) and the substitution rate in substitutions per site per million years is displayed.* ***b)*** *The same analysis as in panel a was performed on RD2. Note that the y-axes in panels a and b are not on the same scale.* ***c)*** *The relationship between the number of substitutions per site (S) and the corresponding time points (T) using RD1. A power-law TDRP model was fitted to the data. Values for the goodness and significance of fit are provided. The fit of a linear model is given for comparative purposes. This is an example of one model fit on a subset of 500 values per calibration point. For TDRP correction, the power-law model fitting was repeated 1000 times per ancestral node to be corrected.* ***d)*** *The same analysis as in panel c was performed on RD2.*

The TDRP in viruses can be empirically described well by a power-law curve (Aiewsakun & Katzourakis, 2016). We fitted a power-law model to the relationship between the number of substitutions per site (*S*) and the evolutionary time scale (*T*) in million years ago (Fig. SF2c and SF2d). We used this model to correct for TDRP by fitting the power-law model on random sub-samples of corresponding *S* and *T* estimates. Subsequently, we extrapolated the model by predicting the *T* values of a node to be corrected using its *S* estimates (leaving out the node to be predicted). This correction could only be performed for clades in which we sampled the posterior distribution of the tmrca. For each node to be corrected, 1000 iterations of sub-sampling and model fitting were performed using the 18 single calibration points. We corrected a total of 23 ancestral nodes (RD1_corr and RD2_corr in Fig. SF3), corresponding to the tmrca of all 18 calibration points, the tmrca of each crown-group, the tmrca of all *Mammals* (with one exception), the tmrca of the *Aves/Testudines* clade, and the tmrca of the split between the *Aves/Testudines* and *Mammals*.

Our results show an overall good fit of the power-law model on the PV data (RD1: *adj.* $R^2$ = 0.912, *min* = 0.904, *max* = 0.920; RD2: *adj.* $R^2$ = 0.930, *min* = 0.924, *max* = 0.935). Interestingly, the data set used for correction seems to have an impact, as in RD2 the corrected node age is always older than the corresponding correction in RD1 (Fig. SF3). This might be due to the lower substitution rates obtained in RD2, as a consequence of differences in representative taxa in the two reduced data sets. When comparing the corrected times using single calibration points (RD1_corr and RD2_corr in Fig. SF3) versus the inferred time using multiple calibration points on RD1 and RD2 (RD1 and RD2 in Fig. SF3), we observe that older nodes are corrected to have a younger age, while younger nodes are corrected to older ages.

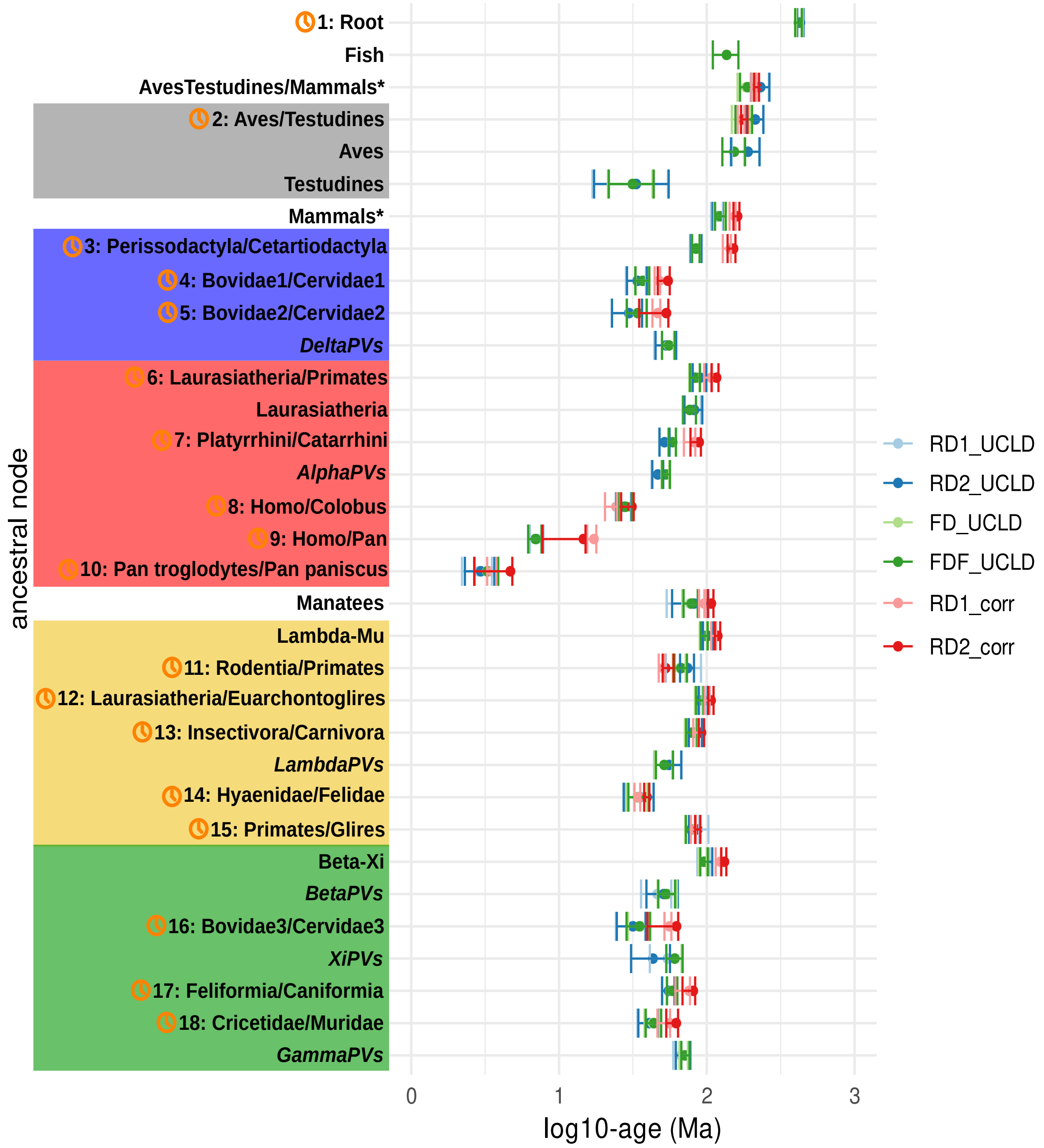


*Figure SF3. Inferred age for ancestral nodes on dated bayesian phylogenetic trees. This figure allows to compare the inferred age on the different data sets: FD, FDF, RD1, and RD2. The phylogenetic trees were inferred by time inference using the uncorrelated relaxed clock model with a Yule tree prior and 18 calibration nodes. The nodes are ancestors of clades within the PV crowngroups and unclassified clades, that are in the same colour code as in Fig. 1 in the main text. The nodes used for calibration are indicated with a clock symbol and the corresponding number. The node age is in million years ago (Ma), and on a log10 scale. For certain nodes, TDRP correction for the age have been made: RD1_corr and RD2_corr. For FD, FDF, RD1, and RD2, error bars encompass 95% HPD for the age of the nodes. For RD1_corr and RD2_corr, error bars encompass the lowest and highest inferred 95% CI for the 18 different nodes used for error correction.*

To obtain a better overview of the differences between using 18 multiple calibration points, 18 single calibration points, and the 18 corrections made for these, we calculated the relative difference as $(T_{multiple} - T_{single}) / T_{single}$ and $(T_{corrected} - T_{single}) / T_{single}$ (Fig. SF4). We observe that for both RD1 and RD2, the differences between multiple and single calibrations are smaller than between corrected and single calibrations. The differences for the corrected calibrations are more important for younger nodes: the younger the node used for calibration, the faster the molecular clock ticks. When fitting a linear model to the relative differences of the multiple and corrected calibrations, we observe that the negative correlation between the corrected calibrations and the host divergence time is significant for nodes between 0 and 100 Ma, in both RD1 and RD2 (Fig. SF4).

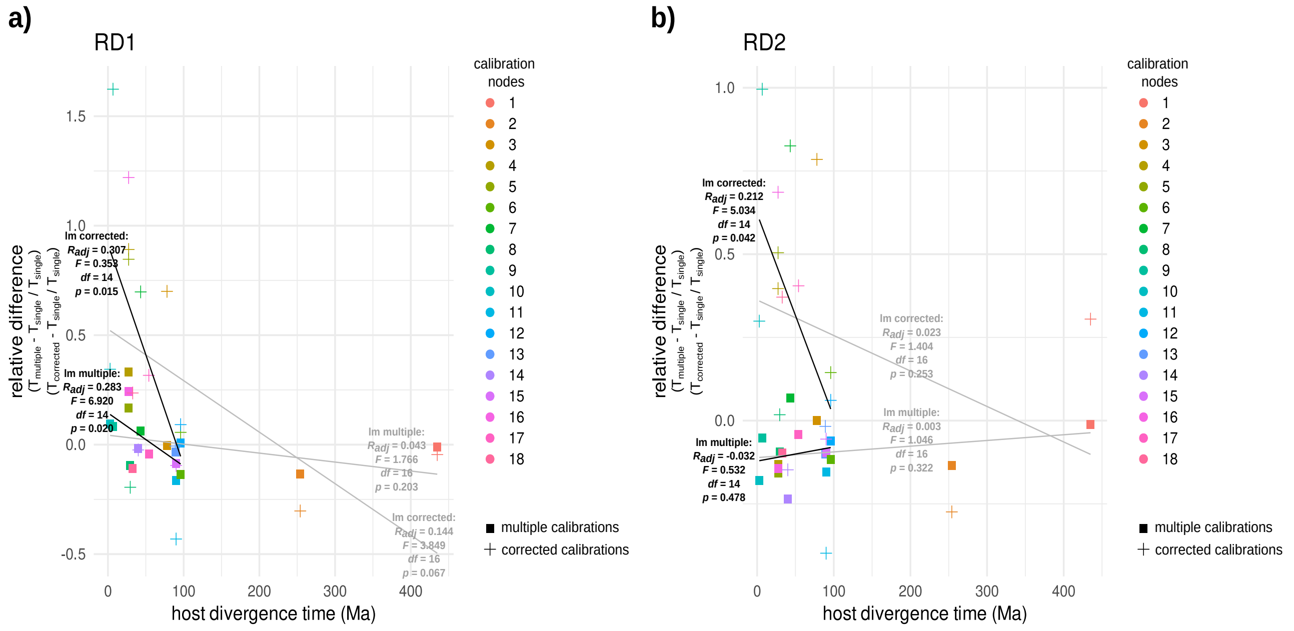


*Figure SF4. Relative differences of the inferred PV divergence times on RD1 and RD2. The differences are shown for the 18 multiple and the 18 corrected calibration points, relative to the 18 single calibration points. The grey lines indicate a linear model fitted to all data points (multiple and corrected) and the black lines indicate a linear model fitted to all young data points with a host divergence time below 100 Ma.*
